# Regulatory and evolutionary impact of DNA methylation in two songbird species and their naturally occurring F_1_ hybrids

**DOI:** 10.1101/2024.01.18.576185

**Authors:** Jesper Boman, Anna Qvarnström, Carina F. Mugal

## Abstract

Regulation of transcription by DNA methylation in 5’-CpG-3’ context is a widespread mechanism allowing differential expression of genetically identical cells to persist throughout development. Consequently, differences in DNA methylation can reinforce variation in gene expression among cells, tissues, populations and species. Despite a surge in studies on DNA methylation, we know little about the importance of DNA methylation in population differentiation and speciation. Here we investigate the regulatory and evolutionary impact of DNA methylation in five tissues of two *Ficedula* flycatcher species and their naturally occurring F_1_ hybrids. We show that the density of CpG in the promoters of genes determines the strength of the association between gene expression and DNA methylation. The impact of DNA methylation on gene expression varies among tissues with brain showing unique patterns. Differentially expressed genes between parental species are predicted by genetic– and methylation differentiation in CpG-rich promoters. However, both these factors fail to predict hybrid misexpression suggesting that promoter mismethylation is not a main determinant of hybrid misexpression in *Ficedula* flycatchers. Using allele-specific methylation estimates in hybrids we also determine the genome-wide contribution of *cis-* and *trans* effects in DNA methylation differentiation. These distinct mechanisms are roughly balanced in all tissues except brain, where *trans* differences predominate. Overall, this study provides insight on the regulatory and evolutionary impact of DNA methylation in songbirds.

## Introduction

Genetic lineages that diverge and form new species acquire both genetic and phenotypic changes in the process. As a by-product, intrinsic reproductive isolation in the form of hybrid dysfunction may evolve. The genetic basis of hybrid dysfunction is usually interactions between incompatible alleles that have never been tested in the same cellular environment [1]. These so-called Bateson-Dobzhansky-Muller incompatibilities (BDMIs) [2–4] are believed to be important for the completion of speciation since they are not dependent on the environment [5]. While some progress have been made in mapping BDMI loci [6], the molecular mechanisms causing hybrid dysfunction is less clear [7]. Incompatible interactions between transcription factors and their binding sites is a class of BDMIs that may cause low fitness in hybrids by the misexpression of genes [8]. Misexpression in hybrids could also be caused by other gene regulatory mechanisms e.g. aberrant levels of different epigenetic marks such as DNA methylation [9]. Given the large number of genes, transcription factors and *cis*-regulatory elements in a vertebrate genome, gene regulatory mechanisms are likely to be an important albeit understudied source of BDMIs [10, 11]. With large genomic data sets we can now both characterize genetic and epigenetic sources of gene regulatory variation and thus gain insight not only on the degree of hybrid misexpression, but also the molecular basis of misregulation [12–14]. In this study we specifically assess the role of DNA methylation in gene misexpression of naturally occurring hybrids.

In vertebrates, DNA methylation is most frequently occurring at cytosines in the 5’-CpG-3’ dinucleotide context and is often associated with transcriptional repression [15]. How DNA methylation regulates gene expression is explained by the “molecular lock” model [16]. Following *de novo* DNA methylation, a gene is locked in a silent state preventing further transcription until the DNA methylation marks are removed, either passively through lack of DNA methylation maintenance during replication or actively using enzymatic activity. In addition to silencing genes, DNA methylation in vertebrates is also used to silence the expression of transposable elements (TEs) [17–19]. Keeping genes (and TEs) repressed through DNA methylation is thought to occur through three main mechanisms that are not mutually exclusive: 1) DNA methylation prevents binding of transcription factors to target sequences [20], 2) DNA methylation acts as substrate for proteins mediating repression such as MeCP2 [21], or 3) *de novo* methylation alters the chromatin structure to a more compact inactive state [22]. Common to these mechanisms is that they are all predicted to yield a negative relationship between the level of DNA methylation in the promoter region and expression level. Gene body methylation, on the other hand shows (if any) a positive or quadratic relationship with expression level, but the function of gene body methylation is debated [23–27].

While most of the genome is methylated in adult animals, some sequence regions escape the wave of *de novo* methylation occurring during development [28]. These so-called CpG islands (CGI) are characterized by a high density of CpG dinucleotides compared to the level predicted by their GC content [29]. Typically, unmethylated CpG dinucleotides contribute to an open chromatin state permissible for transcriptional initiation [30]. Roughly 70 % of all promoters in the human genome have CGIs [31]. In humans, most CpG-dense promoters (CGI promoters) associated with housekeeping genes remain unmethylated in adult tissues, while other CGI promoters are dynamically regulated [32]. In contrast, questions remain to what extent CpG-deficient promoters are regulated by DNA methylation [32, 33].

Differences in methylation caused by genetic changes among alleles, populations and species can arise from two mechanisms: either *cis*-regulatory changes specific to the DNA sequence at a locus or *trans*-regulatory changes because of divergence in structure or regulation of interacting factors with potential to change methylation level. In model organisms, this relationship has been investigated using data from hundreds or thousands of individuals to determine quantitative trait loci affecting methylation level [34]. These regulatory mechanisms may also be distinguished using F_1_ hybrid individuals, which was pioneered by studies of gene expression divergence in mice and fruit flies [35, 36], but also recently applied to other molecular phenotypes [37, 38]. The hybrid represents a *trans environment* in which allele-specific measures of molecular phenotypes can be measured and contrasted with parental species. Here, we extend this approach to DNA methylation data and develop a statistical framework to infer the molecular mechanism of DNA methylation differentiation.

Specifically, in this study, we investigate the regulatory and evolutionary impact of DNA methylation in two *Ficedula* flycatcher species and their naturally occurring F_1_ hybrids. The pied flycatcher (*Ficedula hypoleuca*) and the collared flycatcher (*F. albicollis*) are two species of songbirds that diverged approximately 1 MYA [39]. Interspecific mating occurs in sympatry e.g. on the island of Öland in the Baltic Sea [40]. F_1_ hybrids of both sexes are infertile [41, 42], show reduced viability [43], and males have an increased metabolic rate [44, 45]. A number of studies has investigated DNA methylation in birds [e.g. 46–54]. In general, methylation patterns in birds are similar to other vertebrates. However, compared to mammals much less is known about genome-wide DNA methylation patterns across tissues. Also, the interplay between DNA methylation, gene expression and genetic variation, during speciation and divergence (i.e. the evolutionary impact of DNA methylation), remains largely unexplored. For this reason, we first provide a detailed examination of the association between DNA methylation and gene expression across five different tissues. We then identify differentially methylated regions (DMRs) on a genome-wide scale between tissues, species and between parental species and F_1_ hybrids. We also investigate molecular mechanisms that could explain the differentiation in DNA methylation as well as inheritance patterns of DNA methylation. Finally, we explore the role of genetic vs. epigenetic change in differential gene expression among species as well as in hybrid misexpression.

## Results

### Study system and sequence data

We performed whole-genome bisulfite sequencing from samples of five tissues: brain, heart, kidney, liver, and testis in 14 wild-caught male flycatchers belonging to two *Ficedula* flycatcher species and their naturally occurring F_1_ hybrids: 6 collared flycatchers, 5 pied flycatchers and 3 F_1_ hybrids (41.5 billion reads in total). All F_1_ hybrids (HYB) were offspring from crosses between female pied (PIE) flycatchers and male collared (COL) flycatchers [55]. Each of the 70 samples was sequenced using 2-9 technical replicates, corresponding to 372 technical samples in total (Table S1). On average, 592 M reads were obtained per biological sample. We then collated the whole-genome bisulfite sequencing data with previously sequenced RNA-seq data from the same 70 tissue samples [55], thus forming matched genomic data sets.

### Association between DNA methylation and gene expression across tissues

We assessed the average DNA methylation profile across a set of 8563 genes with annotated 5’ and 3’ UTRs, i.e. defined transcription start sites (TSSs) and transcription termination sites (TTSs). For this purpose, we *a priori* split the genes into two sets based on presence or absence of CGI annotation in their promoters, referred to as CGI and *Other* promoters, respectively. Following Mugal *et al.* [55], we defined promoter regions as the 2 kb upstream region of the TSS. CGI promoters made up 59% of the promoter set. We further split the genes into three categories of gene expression level: low (L: 20% of gene with the lowest gene expression level), high (H: 20% of gene with the highest gene expression level), and medium (M: 60% remaining genes with an intermediate gene expression level).

We computed the DNA methylation profile from 5 kb upstream of the TSS to 5 kb downstream of the TTS separately for tissues and the different sets of genes. Among tissues and irrespective of promoter type, brain showed the highest average methylation level and testis the lowest (Figure 1A-J). On average gene bodies showed higher methylation levels than the 5 kb up– and downstream regions. Splitting the gene body into exons and introns revealed a higher methylation level in exons (Figure S1). Genes with CGI promoters showed a drop in DNA methylation levels especially around the TSS (Figure 1A-E). Genes with low expression generally showed higher promoter methylation, in particular for CGI promoters, but in most tissues lower gene body methylation than the other categories, regardless of promoter type. The exception to the latter pattern was brain which showed the lowest gene body methylation for genes with the highest amount of expression.

**Figure 1.**
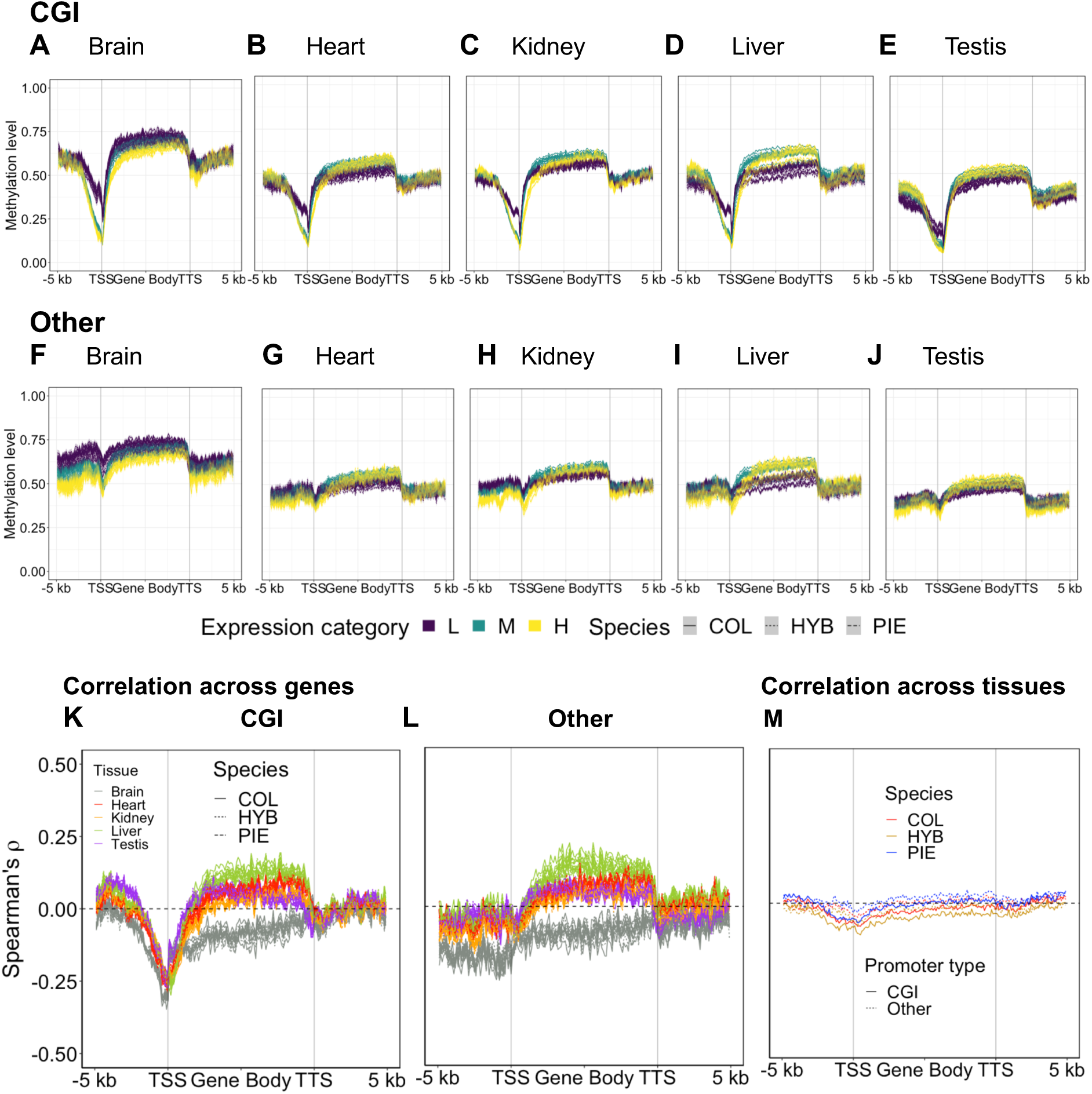
Relationship between DNA methylation and gene expression. Average methylation level was plotted in up– and downstream as well as gene bodies for genes with CGI (A-E) and *Other* (F-J) promoters. Correlation profile between DNA methylation and gene expression across genes (K & L) and across tissue (M).

To further assess the relationship between DNA methylation and gene expression, we computed a gene profile of their correlation across genes (Figure 1K, L). Here, all tissues except brain showed a weak positive correlation between gene body methylation and gene expression regardless of promoter type. For brain, the relationship was negative. Correlating gene expression and DNA methylation across genes also revealed a clear difference between promoter types. While higher methylation levels in CGI promoters were consistent with lower expression (Figure 1K), methylation levels in *Other* promoters only showed a negative relationship with gene expression for brain (Figure 1L).

We also correlated gene expression and DNA methylation for each gene across tissues within groups of samples, which investigates if variation in DNA methylation across tissues is associated with variation in gene expression across tissues. Since DNA methylation is set during tissue differentiation, we take this as a proxy to investigate if gene expression is set during differentiation (Figure 1M). Similar to the correlations among genes within each tissue, DNA methylation at CGI promoters were more negatively correlated with gene expression than in *Other* promoters, but the overall effect was weaker than correlations across genes. A weak negative correlation was observed across tissues for gene body methylation and expression. For these general patterns of gene body and gene-proximal DNA methylation, the difference between COL, HYB and PIE was marginal. To conclude, these results show that promoter type is an important predictor for the association between promoter methylation and gene expression and that CGI promoter methylation is associated with tissue-specific gene expression.

### Tissue-specific patterns of DNA methylation

We identified regions with tissue-specific patterns of DNA methylation on a genome-wide scale. For this purpose, we called differentially methylated regions (DMRs) between tissues using the BSmooth method [56], separately per species (COL, HYB and PIE). We then called regions with tissue-specific methylation (tsDMRs) by identifying DMRs unique for a certain focal tissue that is significantly differentially methylated in comparisons to all other tissues. For all sample groups, testis had the highest number of tsDMRs, followed by heart in COL and brain in PIE and HYB (Figure 2A-E). In heart, kidney, and liver most tsDMRs were hypomethylated while the reverse was true in testis (Table S2). We here use hyper– and hypomethylated as relative terms of lower and higher methylation in a comparison, following Hansen et al. [56]. Our findings therefore indicate that tissue-specific methylation can on average either have permissive or repressive functions, dependent on tissue.

**Figure 2.**
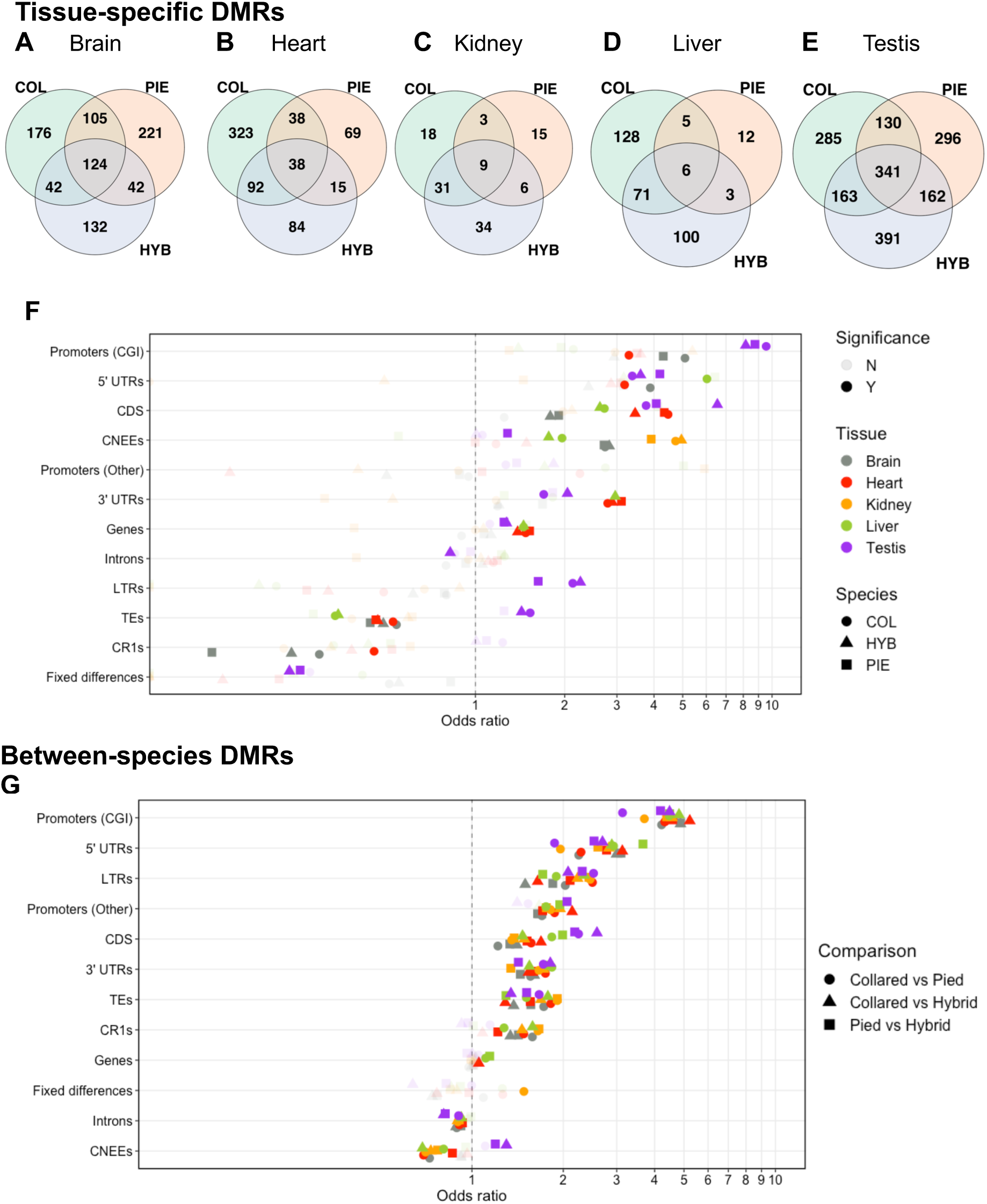
Differentially methylated regions between tissues and species. Tissue-specific DMRs (tDMRs) were shared between sample groups to a much greater extent than expected by chance (<1 in all cases; A-E). Enrichment analysis of tsDMRs and annotation features (F). Enrichment analysis of between-species DMRs (spDMRs) and annotation features (G). Significance in all cases was calculated using 1000 Monte Carlo replicates with a family wise error rate (FWER) of 0.1. In (F-G), “fixed differences” refer to the 100± bp vicinity of fixed differences.

It is possible that functionally important tissue-specific patterns are shared between PIE and COL, and potentially also present in viable F_1_ hybrids. For all tissues, more than half of tsDMRs shared between COL and PIE were also found in HYB (Monte Carlo *p-*value ≈ 0; Figure 2A). This indicates that the identified tsDMRs represent functionally important genomic regions with constrained methylation levels. Next, we investigated the overlap between tsDMRs and regions with functional annotation (Figure 2F). A general pattern emerged with tsDMRs enriched in regulatory regions such as 5’ UTRs and promoter regions (especially CGI promoters). Also, conserved non-exonic elements (CNEEs), which are suggested to function as tissue-specific regulatory elements [57], were enriched in tsDMRs in most tissues. Transposable elements (TEs) were underrepresented in all tissues except in testis, which also showed the greatest bias towards hypermethylation and thus likely repression among their tsDMRs.

We overlapped tsDMRs shared by COL, PIE and HYB with genes and their ±5 kb up– and downstream regions to investigate if tsDMRs show gene ontology (GO) terms consistent with tissue-specific function (Table S3-6). This revealed that brain-specific DMRs were enriched for genes involved in ion transport, while kidney-specific DMRs were enriched in *HOX* genes with developmental function. Heart-specific DMRs showed no significant enrichment while testis-specific DMRs were enriched in e.g. autophagy but also cardiac cell functions in concordance with most of testis-DMRs being hypermethylated (Table S2). These findings highlight that the inferred tsDMRs show signatures of tissue-specific repression as well as permission [58]. We next investigated the relationship between tsDMRs overlapping with genes and tissue-specific expression and found evidence for an association in brain and testis (Supplementary Results 2). One interesting example of tissue-specific regulation is that of *KDM2A*, a gene that represses transcription and is involved in pericentromeric chromatin formation through binding unmethylated DNA [59], possibly an important function in the demethylated germline genome. Three conserved tsDMRs are found in testis inside intron 11 of *KDM2A*. One of these testis-specific DMRs completely overlaps a CGI, which may act as an alternative promoter for a shorter 5’-truncated forms of *KDM2A*, which are known from other species (Turberfield et al. 2019).

### Differential methylation between species is enriched in regulatory regions

We identified DMRs between species (spDMRs) by pairwise comparison between COL, PIE and HYB, in order to investigate general patterns of DNA methylation differences between the two parental species and their viable F1 hybrids. Brain showed the highest number of spDMRs (11330) between species while testis had the lowest (7430) (Table 1). For all tissues, COL were significantly more often hypermethylated compared to PIE (*p*-value < 2.2 x 10^-16^; Binomial Test). In HYB, the pattern varied among tissues, with lower methylation in heart and higher methylation in testis than both parentals. For all tissues, there were more DMRs in the comparison between PIE and HYB than between COL and HYB. This means that F1 hybrids have a methylation level closer to COL than PIE. Since all crosses had a COL sire this could either indicate that the paternal methylation pattern has a greater influence on methylation levels in male offspring than the maternal or collared-dominant inheritance.

**Table 1.**
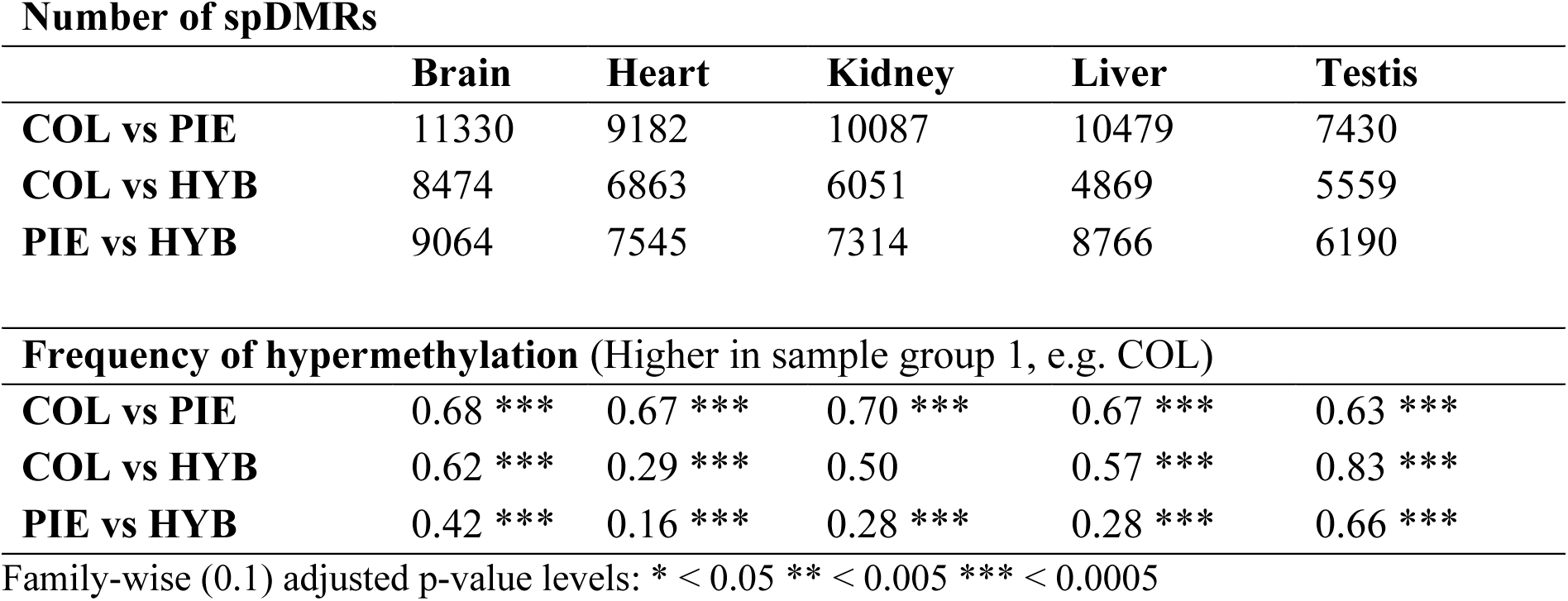
Number of spDMRs and the frequency of hypermethylation per comparison. Values were rounded to two decimal points.

| Number of spDMRs |  |  |  |  |  |
| --- | --- | --- | --- | --- | --- |
|  | Brain | Heart | Kidney | Liver | Testis |
| COL vs PIE | 11330 | 9182 | 10087 | 10479 | 7430 |
| COL vs HYB | 8474 | 6863 | 6051 | 4869 | 5559 |
| PIE vs HYB | 9064 | 7545 | 7314 | 8766 | 6190 |
| Frequency of hypermethylation (Higher in sample group 1, e.g. COL) |  |  |  |  |  |
| COL vs PIE | 0.68 *** | 0.67 *** | 0.70 *** | 0.67 *** | 0.63 *** |
| COL vs HYB | 0.62 *** | 0.29 *** | 0.50 | 0.57 *** | 0.83 *** |
| PIE vs HYB | 0.42 *** | 0.16 *** | 0.28 *** | 0.28 *** | 0.66 *** |

Family-wise (0.1) adjusted p-value levels: \* $< 0.05$ \*\* $< 0.005$ \*\*\* $< 0.0005$

We next investigated the overlap between genome features and spDMRs. This revealed a significant enrichment of spDMRs in regulatory regions such as promoters and UTRs (Figure 2; adjusted Monte Carlo *p*-value < 0.05, for a sample size of 1000). In contrast, other putative regulatory regions such as CNEEs were significantly underrepresented among spDMRS in most tissues except testis, in all pairwise comparisons. This indicates that CpG methylation at CNEEs may be under functional constraint. Transposons, in particular, long terminal repeat (LTR) retrotransposons were enriched in spDMRs.

### DNA methylation patterns varies more among tissues than species

Our results showed that tsDMRs and spDMRs had similar enrichment patterns in promoters and UTRs, but distinct enrichment patterns in CNEEs and TEs (Figure 2). In order to assess the relative contribution of tissue and species to methylation variation within annotation sets quantitatively, we performed between-groups principal component analysis (BCA), and tested the significance of the amount of explained variation (*R^2^*) using permutations [60]. Overall tissue differences in methylation level were much greater than evolutionary difference between the species (HYB excluded; Table 2), where TEs showed the lowest between-tissue *R^2^* (51%) and highest *R^2^* between species (11%) among annotated features. The ±100 bp region around fixed differences showed the greatest between-species effect (16%), indicating a *cis-*genetic coupling between genetic and DNA methylation differences. Annotations with higher between-tissue *R^2^* had in general lower between-species *R^2^* even after controlling for tissue.

**Table 2.** Between-groups principal component analysis. *R^2^* is the proportion of variance explained. Values were rounded to two decimal points.

| Annotation | $R^2_{\text{between tissues}}$ | $R^2_{\text{between species}}$ | $R^2_{\text{between species controlled for tissue}}$ |
| --- | --- | --- | --- |
| Promoters (CGI) | 0.75 * | 0.03 | 0.12 * |
| Promoters (*Other*) | 0.65 ** | 0.05 | 0.14 * |
| Genes | 0.88 * | 0.02 | 0.13 * |
| Introns | 0.55 * | 0.09 * | 0.20 * |
| CNEEs | 0.61 * | 0.04 | 0.11 * |
| CDSs | 0.67 * | 0.04 | 0.11 * |
| 5' UTRs | 0.68 * | 0.04 | 0.13 ** |
| 3' UTRs | 0.64 * | 0.05 | 0.15 * |
| TEs | 0.51 * | 0.11 * | 0.23 * |
| CR1s | 0.49 * | 0.12 * | 0.24 * |
| LTRs | 0.55 * | 0.10 * | 0.22 * |
| Fixed differences | 0.47 * | 0.16 * | 0.30 * |

Family-wise (0.1) adjusted $p$ -value levels: \* $< 0.05$ \*\* $< 0.005$ \*\*\* $< 0.0005$

### Genetic differentiation is correlated with differences in methylation between species

We tested the association between absolute methylation differentiation (*M_diff_*) and genetic differentiation (*F_ST_*) between COL and PIE across genes (Figure 3A-C). In general, we observed a clear positive correlation between *F_ST_* and *M_diff_* of ∼0.15-0.25 for *F_ST_* based on CpG sites in the reconstructed ancestor (CpG *F_ST_*) and ∼0.05-0.1 for *F_ST_* based on other sites (non-CpG *F_ST_*) (Figure 3D). There was some variation between tissues, with a stronger correlation coefficient in kidney and brain and lowest in liver (Figure S2). For CGI promoters we observed a strong reduction in correlation in the promoter region using both CpG *F_ST_* and non-CpG *F_ST_*. This region is close to the peak of annotated CGIs along the gene profile (Figure S3). Assuming a causal effect of genetic change on DNA methylation differences among species, this could mean that the DNA methylation level at CGI promoters is under purifying selection and that only genetic changes that do not disrupt methylation level are tolerated to segregate at appreciable frequencies. In support of this hypothesis, we do not observe this reduction in correlation for genes with *Other* promoters, which we previously determined had a weaker relationship with gene expression level and thus epigenetic changes may have less of an effect on expression (Figure 1L). Furthermore, *M_diff_* is three times lower in the promoter region of CGI promoters compared to *Other* promoters (Figure 3A). Both non-CpG *F_ST_* and CpG *F_ST_* are higher in promoters compared to the gene body (Figure 3B-C), especially the exons, which show lower *F_ST_* compatible with purifying selection [61]. This indicates that genetic differentiation in promoter regions is on average tolerated in both CGI and *Other* promoters but less so when affecting CpG sites in CGI promoters.

**Figure 3.**
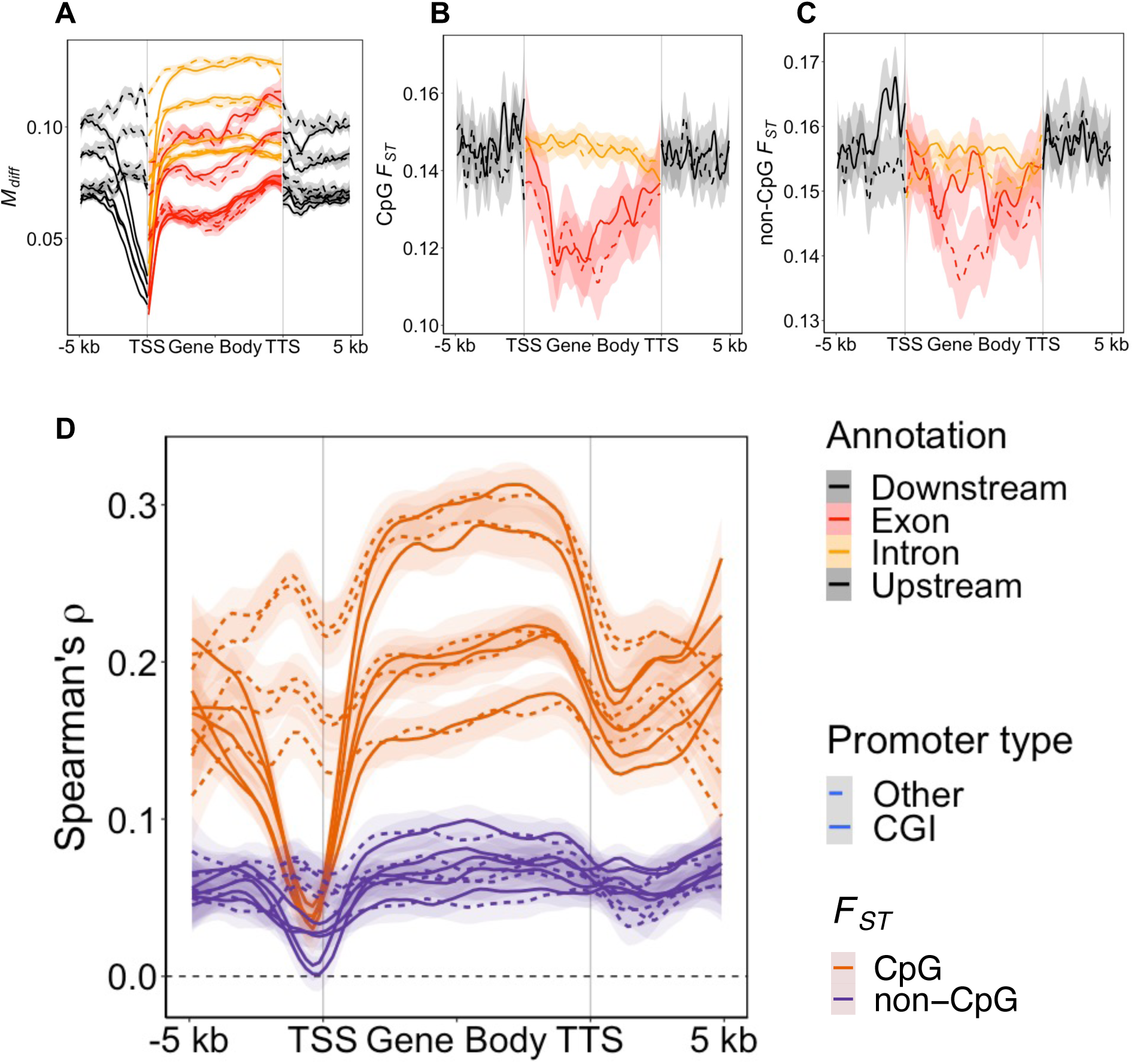
Association between genetic– and methylation differentiation. Correlation between *F_ST_* and *M_diff_* (A). Gene profile for *M_diff_* (B). Gene profile for non-CpG (C) and CpG *F_ST_* (D). Different lines in (A) and (B) are different tissues. Different line types represent the average for different promoter types. See Fig. S2 for (A) and (D) colored by tissue. Lines in plots represent loess curves with shaded region representing the 95 % confidence interval of the local regression.

### Molecular mechanism of DNA methylation differentiation

Changes in DNA methylation that are caused by genetic differentiation between species can either be due to substitutions that affect the local genomic region (*cis*) or elsewhere in the genome (*trans*). These molecular mechanism may be distinguished through contrasting allele-specific effects in F1 hybrids with parental differences [36]. For this purpose, we developed a statistical framework applicable to DNA methylation data inspired by previous categorization systems for gene expression [36, 62]. We classified loci into categories based on differences in DNA methylation between the two parental species as well as between the two alleles in the HYB using a beta regression model and an FDR of 0.1 (Table S7). To distinguish between alleles in the hybrid, we polarized bisulfite-seq reads based on fixed differences between COL and PIE, and calculated allele-specific methylation. In total around 37k fixed differences per tissue were used as markers (Table S8). We measured the allele-specific methylation in the ±100 bp region from a fixed difference. Most loci either had no CpG site or too low coverage to be considered. This restricted the dataset to 3251, 5939, 6120 and 7701 loci in brain, heart, liver and testis respectively.

In our analysis we focused on the relative contribution of *cis* and *trans* effects on DNA methylation and if this varies with tissues and sequence divergence. A prediction based on gene expression differences in flies and yeast is that the proportion of *cis*-changes should increase with sequence divergence [63, 64]. Since the Z sex chromosome is more genetically differentiated between flycatchers than autosomes [65], we performed the analysis separately for Z and autosomes. In total, 1.6% of all callable loci across tissue showed a statistically detectable difference in at least one of the pair-wise comparisons COL/PIE or the two alleles in the HYB (Figure 4A). Distribution of regulatory patterns were significantly different among tissues (Figure 4; Fisher’s exact test, *p ≈* 0.0002). Brain showed an excess of conserved loci (Figure 4B). Among divergent (non-conserved) loci there was also a difference between tissues (*p* ≈ 0.01). An excess of *trans* differences was found on brain autosomes (Binomial test, *p* ≈ 0.001). For other tissues and chromosome types, an even contribution of *cis* and *trans* differences could not be rejected. Overall, distribution of divergence categories was not significantly different between autosomes and the Z chromosome (*p* > 0.05). The exception was testis (*p* ≈ 0.001), which showed an excess of non-conserved loci on the Z compared to autosomes. A trend with a larger proportion of *cis* relative to *trans* on Z compared to autosomes were observed (Figure 4B) but was not significant for any tissue (Fisher’s exact test, *p >* 0.05).

**Figure 4.**
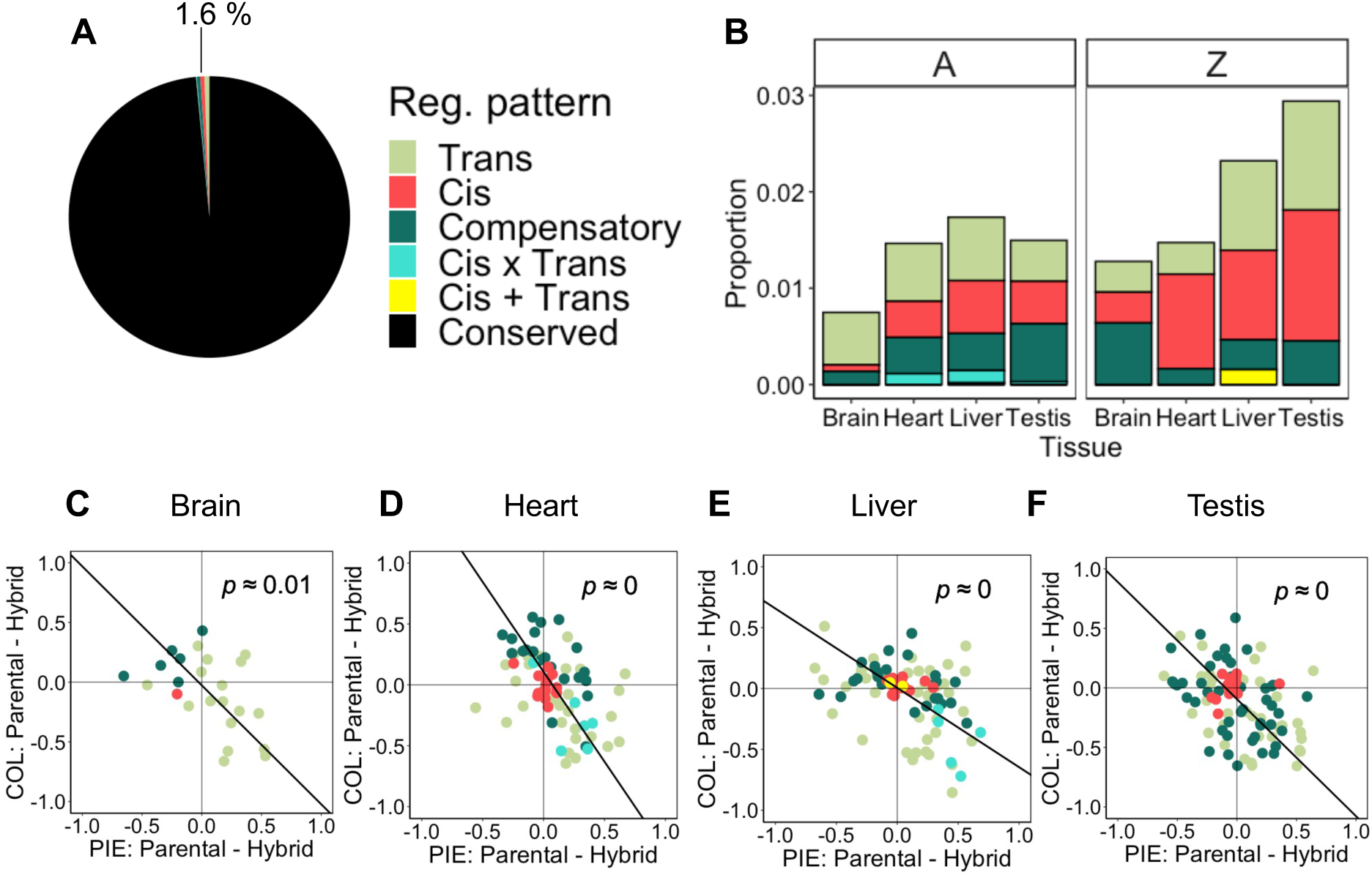
Molecular mechanism of DNA methylation differentiation. (A) Proportion of different classes of molecular mechanisms of DNA methylation differentiation at callable loci. (B) Proportion of divergent (non-conserved) classes per tissue and chromosome type. (C-F) Difference between COL and PIE alleles in their native parental and hybrid cellular environment. Loci with *cis*-only effects are clustered close to the origin as predicted when the methylation level is independent of cellular environment. In contrast, loci with *trans* effects show greater variation but the general trend is a spread along the additive inheritance axis. Bold black lines in C-F are major axis regression lines. Values of *p* in C-F are rounded to two decimals.

We next determined the inheritance pattern of methylation at these loci and investigated its relationship with allele-specific methylation in HYB. This revealed that the inheritance pattern is tightly related to the mechanism of molecular divergence (Figure 4C-F). Major axis regressions between parental and hybrid differences were negative for all tissues (*p* < 0.05), indicating an overall pattern of additivity (Figure 4C-F). Subtle differences in spread are visible among tissues, with liver having the largest variation between PIE in parental and in HYB indicating more COL dominance. The results illustrate two different mechanisms of additive inheritance of a molecular phenotype. Loci with methylation difference due only to *cis* effects showed essentially no difference in methylation level whether in parentals or in hybrids which indicate that they are strictly additively inherited. In other words, *cis* loci are unaffected by the HYB cellular environment. Other loci are affected by the HYB cellular environment unveiling a *trans* effect. That *trans* difference here generally makes the alleles in HYB more similar to each other which creates a different route to additive inheritance since the overall state is in-between parentals. These two types of additive inheritance are only distinguishable when measuring allele-specific methylation.

### Tissue-specific association of differential expression with genetic– and epigenetic change between collared– and pied flycatchers

Much of the interest in DNA methylation lies in its ability to regulate gene expression. Nevertheless, it is not clear to what extent changes in DNA methylation are involved in the evolution of differential expression (DE) between species. Besides DNA methylation patterns, also genetic changes in promoter regions are expected to affect transcriptional regulation and could ultimately result in DE. We sought to test the relative importance of changes in promoter methylation and genetic change in the evolution of DE between COL and PIE. Here, changes in promoter methylation are used to assess epigenetic change. We used non-CpG *F_ST_* to assess genetic changes to more clearly separate genetic and epigenetic effects. First, we explored the patterns of differentiation around well-annotated genes (having both 5’ and 3’ UTR). The number of differentially expressed (FDR=0.1) genes between COL and PIE were lowest in brain and highest in testis (Table S9). As reported above (Figure 2C), between-species DMRs are enriched in the CGI promoter region, with an elevated enrichment in DE compared to non-DE genes (Table 3, Figure 5A). Notably, this same effect is not observed in *Other* promoters (Table3, Figure 5C). In addition, we observed no clear difference in DMR frequency in the bodies of DE vs. non-DE genes for most tissues. Genetic differentiation (non-CpG *F_ST_*) is higher in CGI promoters though only significant in brain and testis after multiple-test correction (Table 3, Figure 5B). All tissue and promoter type combinations had significantly higher non-CpG *F_ST_* in the bodies of DE vs. non-DE genes (Table 3, Figure 5B & 5D), revealing a strong association between local genetic differentiation and DE in this system.

**Figure 5.**
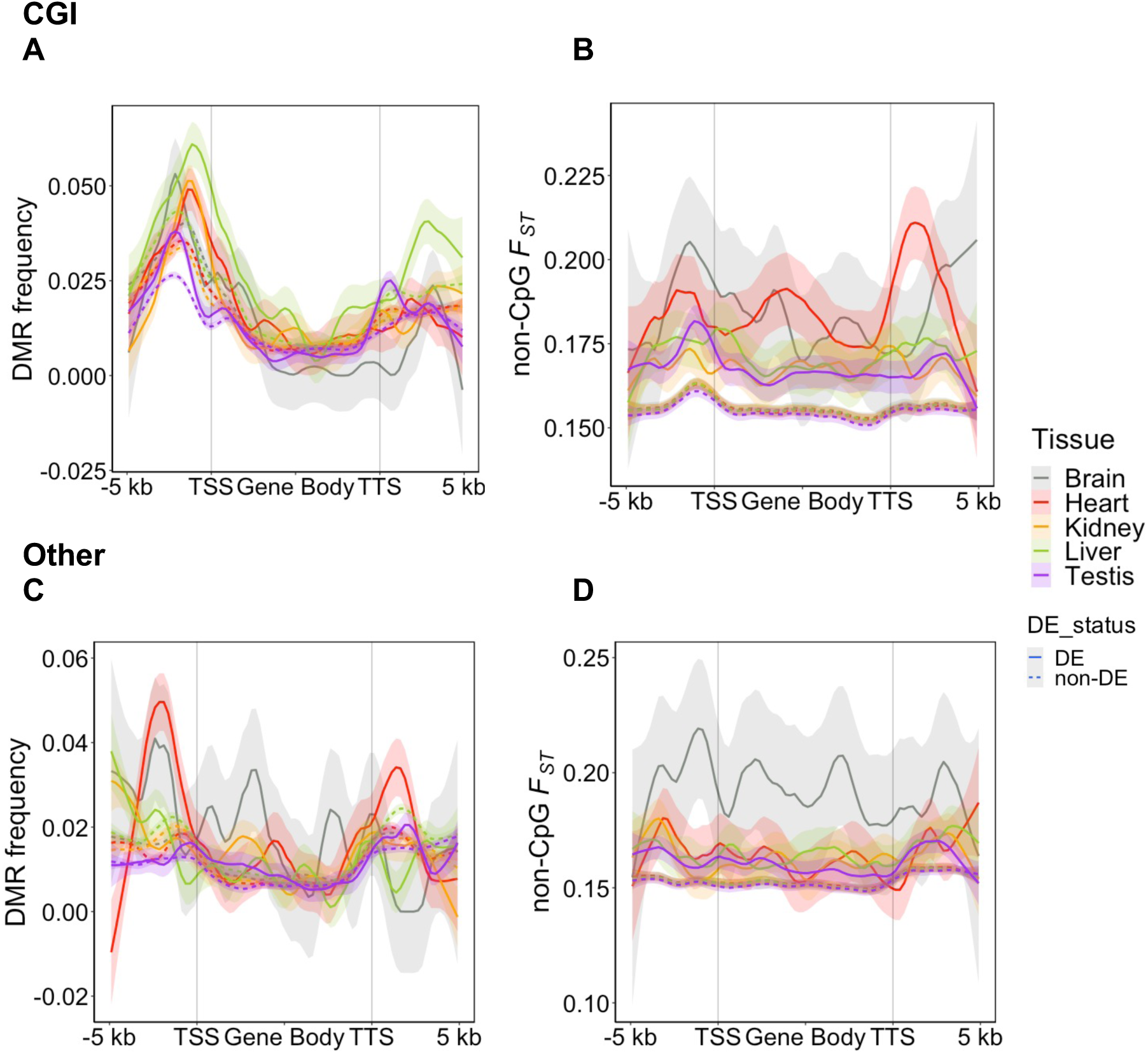
Patterns of genetic and epigenetic change at DE genes between COL and PIE. DE genes had more DMRs in CGI promoters (A) but not in the *Other* promoters (C). DE genes also had higher non-CpG *F_ST_* (B) across the gene bodies in all tissues and for both promoter types (B and D). Lines in plots represent loess curves and shaded regions around lines are the 95 % confidence intervals.

**Table 3.** Determinants in *cis* of DE between COL and PIE. Average DMR frequency differences and non-CpG *F_ST_* in the 2k upstream promoter region and throughout the gene body were compared between DE and non-DE genes using paired *t-*tests. The table displays p-values of those tests.

| Tissue | Promoter type | P<sub>DMR</sub> freq. | GB<sub>DMR</sub> freq. | P<sub>non-CpG</sub> $F_{ST}$ | GB<sub>non-CpG</sub> $F_{ST}$ |
| --- | --- | --- | --- | --- | --- |
| Brain | CGI | 1 | 0.456 (-) | 0.041 * | 0 *** |
| Heart | CGI | 0.023 * | 0.039 * | 0.1 | 0 *** |
| Kidney | CGI | 0.006 * | 1 | 1 | 0 *** |
| Liver | CGI | 0.003 ** | 0 *** | 0.125 | 0 *** |
| Testis | CGI | 0.367 | 1 (-) | 0.001 ** | 0 *** |
| Brain | Other | 1 (-) | 0.069 | 0.013 * | 0 *** |
| Heart | Other | 0.660 | 0.2 | 1 | 0 *** |
| Kidney | Other | 0.171 (-) | 0.023 * | 1 (-) | 0 *** |
| Liver | Other | 0.020 (-) | 1 | 1 | 0 *** |
| Testis | Other | 1 | 1 | 0.456 | 0 *** |

(-) = Average value lower in DE genes
P = Promoter
GB = Gene body
Family-wise (0.1) adjusted p-value levels: \* < 0.05 \*\* < 0.005 \*\*\* < 0.0005

### *Cis*-genetic and –epigenetic changes do not predict misexpression in hybrids

We also investigated whether genetic differentiation between COL and PIE as well as hybrid-specific methylation changes could predict misexpression in hybrids, i.e. DE between HYB and both parental species in the same direction. In other words, an overdominant or underdominant expression pattern. Overall inheritance of methylation patterns for both promoter types were mainly additive (Supplementary Results 3), in concordance with results from methylation differentiation around fixed differences (Figure 4). We observed less mismethylation in CGI promoters compared to *Other* promoters (Supplementary Results 3). In general, neither DMRs between each parental species and HYB nor non-CpG *F_ST_* in promoter regions were higher for DE genes between HYB and both parental species (Figure S4). In gene bodies, non-CpG *F_ST_* was still higher in misexpressed genes in some cases, while significantly lower in heart (Table S10). These results indicate that both *cis-*genetic and *cis-*methylation differences play a relatively minor part in F1 hybrid misexpression patterns and are perhaps dwarfed in importance by *trans* effects or distal *cis* effects.

## Discussion

In this study we investigated both the regulatory and evolutionary impact of DNA methylation in two species of *Ficedula* flycatchers and their F1 hybrids. Using these two layers of analyses we could evaluate both the functional impact of DNA methylation in two wild bird species and its relation to genetic divergence and differential gene expression. In addition, by using F1 hybrids we could investigate the role of methylation in hybrid misexpression, thus probing deeper into the regulatory underpinnings of hybrid dysfunction.

Genome-wide we observed much greater differentiation of DNA methylation between tissues than species as expected. A similar pattern was observed in a study of three primates including humans [66]. Among the tissues studied here, brain showed the most unique methylation profile. Only brain tissue had a negative correlation between gene expression and gene body methylation. This distinctive pattern could be driven by the special role of the *MeCP2* gene, which is expressed at remarkable levels of more than >16M proteins per nucleus in nerve cells of mammals [67]. MeCP2 binds to methylated C’s and recruits the NCoR1/2 co-repressor complex causing transcriptional repression possibly through deacetylating histone tails [21, 68]. This mechanism could potentially make DNA methylation marks at gene bodies a target of chromatin compaction, which could explain why gene body methylation was negatively correlated with expression in this case.

We observed that the relationship between DNA methylation at promoters and gene expression differed among promoters with or without CGIs, which we suggest affect their different evolutionary constraints. DNA methylation level at CGI promoters showed a stronger relationship with gene expression, in congruence with a recent study of fibroblasts from six mammals and chicken [54]. We cannot rule out that some genes with *Other* promoters are also regulated by DNA methylation as have been observed in some systems [23, 33]. Our study highlights the benefits of whole-genome bisulfite sequencing for gaining a complete view of the importance of DNA methylation, in contrast to the popular reduced-representation bisulfite sequencing method which may bias analysis to CpG-rich genes and promoters where variation in DNA methylation may be more impactful [69, 70]. For example, we observed that CGI promoter methylation to a higher degree was associated with tissue-specific expression compared to methylation patterns at *Other* promoters, a conclusion which would be difficult to draw using reduced-representation data generated with CG-specific digestion enzymes.

Beyond promoters and gene bodies, less is known of the regulatory impact of differential methylation. What is for example the impact of DNA methylation levels on enhancer or other kinds of *cis-*regulatory sequences? We observed an enrichment of tsDMRs and a lack of spDMRs in CNEEs, a subset of which putatively acts as tissue-specific *cis-*regulatory elements. DNA methylation at *cis*-regulatory elements impacts binding affinities both positively and negatively, but the analysis is complicated by the fact that transcription factor binding itself seems to induce active demethylation possibly through attracting TET demethylases [20, 71]. Still, we can conclude that flycatcher CNEEs are likely to be involved in tissue-specific regulation of expression potentially through tissue-specific methylation, though experimental studies would be needed for robust confirmation.

While DNA methylation was constrained among species at CNEEs the reverse was observed for TEs. They showed of a lack of tsDMRs and an enrichment of spDMRs. More pronounced interspecific differences in DNA methylation at TEs than at CNEEs have previously been observed in primates [72]. TEs could be showing relatively more variation among species due to a higher turnover of CpG sites resulting from high methylation levels or relaxed purifying selection on methylation level. In addition, active copies of LTR retrotransposons carry regulatory elements which could potentially have a functional role in regulating gene expression [73]. TEs may affect divergence and speciation of lineages in more ways than through their associated regulatory elements affecting expression levels of proximal genes. In hybrids, TEs can be misregulated, which could lead to hybrid dysfunctions or sterility as has been observed for the *P-*element in *Drosophila* [74]. While beyond the scope of this paper, the presented dataset could be used to test this hypothesis. With that said, in flycatchers it is possible that genic effects are causing hybrid male sterility since dysregulation of meiotic genes have been observed through single-cell RNA sequencing of male testes [75].

Despite having a relationship with gene expression and thus stronger functional constraints, we did observe an enrichment of spDMRs in CGI promoters. When examining the gene profile of DMR frequency, we find that the peak enrichment of spDMRs in the promoter is in most cases distal to the TSS within the promoter. Therefore, the enrichment of spDMRs in promoters is not coinciding with the minima of correlation between *F_ST_* and *M_diff_* which is more proximal to the TSS, rich in CGI annotation (c.f. Figure 5A, Figure 3D and Figure S3). Nevertheless, since we observed an enrichment of spDMRs within CGI promoters but a marked drop in *cis*-genetic correlation it is likely that some spDMRs at CGI promoters in this system are driven by *trans* differences in methylation. A non-exclusive explanation would be that DMRs are concentrated to the shores of CGIs, which have been shown to be especially prone to differential methylation [76]. While we could survey genome-wide patterns of *cis* and *trans* differences in methylation using fixed differences as marker loci, we lack power to distinguish their relative contribution to divergence at promoters. Ideally, the F1 hybrid method of determining the molecular mechanism of DNA methylation divergence, is complemented by sequencing of trios (parents and hybrid offspring), which would enable much more and less biased marker loci to be used. Even so, we did observe a stronger species-effect in DNA methylation at fixed differences compared to other annotation sets further indicating that genetic and DNA methylation change are coupled [34]. In spite of this, tissue was a stronger determinant of DNA methylation level than species also at fixed differences, highlighting the importance of studying several organs/tissues or cell types for understanding the evolution of DNA methylation [77].

For loci with fixed sequence differences between the two species, we observed an equal amount of methylation differentiation attributable to *cis* or *trans* effects for all tissues except brain where *trans* differences dominated. In a cross between red-jungle fowl and domestic white leghorn chickens a greater share of *cis* differences was observed in a QTL analysis of hypothalamus tissue [51]. This could be due to tissue– or species-specific patterns or the ∼3x greater genetic differentiation between red-jungle fowl and white leghorn compared to COL and PIE. A greater share of *cis* difference in gene expression with greater sequence divergence has been observed in fruit flies and yeast [63, 64]. Though insignificant, we here observed the same trend for DNA methylation in the contrast between Z and A chromosomes, where the more differentiated Z has a trend of higher share of *cis.* By contrasting the allele effect in HYB with parental species we also observed that *trans* differences generally where additively inherited but through a distinct mechanism where both HYB alleles converged to the midparental value of DNA methylation. This could for example be caused by the relatively unexplored molecular process of transvection [78, 79], in which alleles affect each other’s phenotype, which have previously been observed to affect DNA methylation patterns during meiosis in mice [78].

In the last two decades many studies have presented evidence of misexpression in F1 hybrids of a wide range of species [14, 55, 80–82]. While some patterns are emerging, the regulatory mechanisms underlying hybrid misexpression remains relatively obscure. Dependent on tissue, both higher DMR frequency and larger *F_ST_* in promoters were associated with DE between COL and PIE, while the same variables failed to predict DE between parentals and hybrids. If hybrid misexpression was driven by evolution at proximal *cis-*regulatory elements of many genes we would have expected to find similar patterns in both COL-PIE and parental-HYB comparisons. It has been suggested that incompatibilities between divergent *cis-*regulatory elements and *trans-*acting factors in hybrids result in misregulation of genes [11, 13]. Misregulation evolve quickest when interacting *cis-* and *trans* factors diverge under positive selection [8], which may be the case for a subset of the genes differentially expressed between COL and PIE [83]. However, expression is likely in many cases to be under stabilizing selection [84, 85], which may also cause BDMIs if different compensatory mutations fix in diverging lineages [80]. We did observe compensatory evolution of DNA methylation at fixed differences, though most were additively inherited and consequently not mismethylated in hybrids. In addition, we did not observe greater frequencies of DMRs at promoters of misexpressed DE genes which would be expected in a model where hybrid over– or underdominance in gene expression is caused by loss or gain of promoter methylation. One caveat is that such mismethylation might be so deleterious that we miss it when sampling adult yearlings that survived migration to Africa and back, and thus are likely to be in better condition than the average natural F1 hybrid hatched [43]. In theory, genetic or epigenetic misregulation at a single or few two-locus interactions could potentially cascade throughout the expression network and cause hybrid misexpression at many genes. Misregulation of upstream *cis*-*trans* interactions could overshadow the *cis* effects of methylation and genetic difference associated with DE in the parental comparison. In this model, most misexpressed genes do not have incompatibilities in their promoter regions, instead they are symptoms of a few rare cascading interactions. It is possible that the hybrid misexpression observed here are metabolic responses possibly related to the transgressive metabolic rate observed in F1 hybrid flycatchers [44, 45] and somewhat in line with results in a copepod with known mitochrondrial dysfunction in F2 hybrids [86].

Our results showed that there was a weak but pervasive correlation between non-CpG genetic differentiation and DNA methylation except proximal to the TSS (for CGI promoters). While it is possible that a genetic change affects e.g. the binding of a transcription factor leading indirectly to a change in methylation, it is harder to conceive of a molecular mechanism supporting the other direction of causation though it cannot be entirely disregarded. Recent empirical and theoretical studies suggest that epigenetic variation could act as a first substrate for divergent selection and promote speciation through transgenerational plasticity [87–89]. In birds, epigenetic effects independent of genetic effects is likely to have a limited impact on speciation processes due to weak evidence for genomic imprinting [90, 91], but more research is needed. However, a limited transgenerational inheritance does not mean that epigenetic mechanisms are unimportant for speciation in birds and other vertebrates. Epigenetic mechanisms are fundamental in conserving and reshaping transcriptional states, as illustrated by the prominent role of DNA methylation in eye degeneration of the cave morph of the Mexican tetra [92]. In other words, epigenetic mechanisms can play important parts without being the ultimate cause in cases of adaptive differentiation and hybrid dysfunction, both of which are important aspects of speciation.

## Materials and methods

### Sampling scheme and tissue collection

Male flycatchers were collected at the Baltic Island of Öland (57°10’N, 16°56’E) during the breeding season of 2014 [55]. Collected samples included six collared (*Ficedula albicollis*), five pied (*F. hypoleuca*) and three F1 hybrid flycatchers (♀PIE x ♂COL). Tissue collection is described in more detail in Mugal et al. (2020). All sampling procedures were approved by the Swedish Board of Agriculture (Jordbruksverket – DNR 21-11). The following organs/tissues were used for RNA sequencing [55] and whole-genome bisulfite sequencing (WGBS; within the present study): brain (caudal region of the telencephalon) heart, kidney, liver and testis. In addition, we included RNA sequencing data for spleen for the collared flycatcher individuals (Craig et al. 2018).

### Nucleic acid extractions and sequencing

Samples were homogenized using a bead beater with ceramic beads and aliquots were used to extract DNA and RNA. For details on RNA extraction and sequencing, see Mugal et al. (2020). Details on the WGBS are presented in the Supplementary Methods. Overall, 41.6 billion reads were obtained with on average 594 MB reads per biological sample. One kidney sample from one F1 hybrid flycatcher (HYB02) was discarded due to allelic imbalance (data not shown).

### Processing of bisulfite sequence reads and methylation calls

Quality control, filtering– and mapping of reads as well as methylation calls were performed using the reproducible Nextflow workflow v20.10.0 nf-core Methylseq v1.5 [94]. Fixed differences between collared and pied flycatchers – determined using 19 individuals of each species previously sampled on Öland [95] – were masked prior to read-mapping. Bismark v0.22.3 was then used to deduplicate alignments and extract methylation calls for CpG sites. After read mapping and deduplication, the median coverage ranged from 7x to 39x. See Supplementary Methods for more details.

### Methylation level

Since the tissue samples we used in this study consisted of a population of cells each with a possibility of either having methyl mark or not at a certain CpG position, we assessed the methylation status at each CpG position as the proportion of methylated reads. To measure methylation level, we summarized the number of methylated *x_mCpG_* reads mapping to both strands of a reference CpG dinucleotide and divided by the total number of reads *x_uCpG_* + *x_mCpG_*. As regional measure of methylation level covering *n* CpG dinucleotides, we used the average proportion across the individual dinucleotides (*i*),

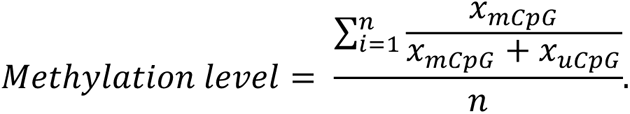

We only included dinucleotides with at least 6– and at most 200 mapped reads unless otherwise stated. The lower limit is included to reduce bias in estimating methylation level caused by lack of data. The upper limit is included to reduce bias induced by collapsed repetitive regions. Other methods and filtering thresholds were used for the smoothed methylation values produced by BSmooth for DMR analysis and BiSeq for the analysis of molecular mechanism of DNA methylation divergence respectively (see below).

### Gene annotation update

With a main goal of improving the annotation of transcription start sites (TSS) and transcription termination sites (TTS) of genes by annotation of their untranslated regions (UTR) we updated the gene annotation for the collared flycatcher genome assembly based on a *de novo* transcriptome assembly. To build the transcriptome assembly, we used RNA-seq reads across all five tissues plus spleen from the six collared flycatchers described above (in total 36 samples). We concatenated all samples and generated four separate and one consensus *de novo* transcriptome assembly using the Oyster-River protocol v2.3.1 [96]. For building gene models, we used MAKER v2.31.10 [97]. Details on transcriptome assembly and gene annotation are provided in the Supplementary Methods.

### RNA sequence read analysis

RNA-seq reads were mapped to the collared flycatcher reference genome, FicAlb1.5 [98] and the updated gene annotation with fixed differences between collared and pied flycatchers masked. All steps up from quality control, read mapping to differential expression analysis were performed using Nextflow v21.02.0.edge nf-core rnaseq v3.0 pipeline [94]. See Supplementary Methods for more details on the pipeline. Differential expression analysis was done with DeSeq2 v1.28 [99] with Salmon count data imported using tximport [100]. We considered genes with an FDR-adjusted *p-*value <0.1 as differentially expressed.

### Additional annotation tracks

We defined promoters as the 2 kb upstream region of the TSS for the set of 9,597genes with at least one transcript with annotated 5’ UTR. CpG islands (CGIs) were inferred using CpG_CLUSTER_ v1.0, with default parameter settings and a minimum length of at least 50 bp [101]. Promoters intersecting CGIs were classified as CGI promoters using BEDtools v2.29.2 [102]. Promoters without any overlapping CGIs were defined as having other types of promoters (*Other*). Phylogenetic conserved elements (CEs) based on PhastCons [103] and a whole-genome alignment of 23 sauropsids were retrieved from (Craig et al. 2018). Conserved non-exonic elements (CNEEs) were defined as CEs which did not overlap exons.

### Gene profile

We investigated the average methylation patterns at genes and their upstream regions (gene profile). To construct a gene profile, we filtered for genes having at least one transcript with 5’-and one with a 3’UTR such that TSS and TTS are defined. Upstream and downstream 5 kb windows of each gene were split into 100 bp nonoverlapping segments. Genes were also split into 100 bp segments, then averaged per 99 ranks across the gene length. Variables of interest, such as methylation level, were averaged across genes in each of these 100 bp segments. For correlation gene profiles, we calculated the Spearman rank correlation coefficient between two selected variables in a segment across all genes using the *cor.test* function in R v4.0.4 [104].

### Identification of differentially methylated regions

Differentially methylated regions (DMRs) between samples were identified using the BSmooth method [56]. We defined DMRs as regions in the 0.01 and 0.99 quantiles of methylation difference, keeping only CpG sites where at least two samples per group (of the pair-wise comparison) have a coverage ≥2. In addition, DMRs needed to span at least 3 CpGs with a mean methylation difference equal to or larger than 0.1. Significant deviation from random (none of the compared groups had more DMRs with higher methylation, i.e. the random expectation is 0.5) in the either direction (hypo– or hypermethylation) for a set of DMRs was determined using binomial tests in R v4.0.4 [104].

### Classification of DMRs

We called DMRs between both tissues and species (spDMRs). We defined tissue-specific DMRs (tsDMRs) as regions with differential methylation in a focal tissue compared to all other tissues using BEDtools v2.29.2 *intersect* requiring at least 25 % reciprocal overlap [66]. Furthermore, to fulfill the criterion of tissue-specificity, the same region was requested not be classified as DMR in any other tissue comparison.

### A method for enrichment analysis between two sets of genomic ranges using resampling

Enrichment analyses of various classes of DMRs in genomic annotation tracks were performed using a custom Bash script employing BEDtools v2.29.2 *intersect* and calculation of empirical *p*-values using a Monte Carlo randomization procedure. For a certain overlap between an annotation track (e.g. introns) and a set of DMRs (e.g. between COL and PIE heart samples), we shuffled the DMRs 1000 times across the genome and calculated the total number of basepairs that overlapped per resampling replicate. To calculate the empirical *p-*value, we compared the overlap in the resampling replicates with that of the actual data and calculated the *p-*value as *r*/*n*, where *r* is the number of replicates with an overlap greater than or equal to the overlap for actual data [105]. *P-*values were corrected using the Bonferroni method to a family-wise error rate of 0.1. Enrichment was defined as the following odds ratio:

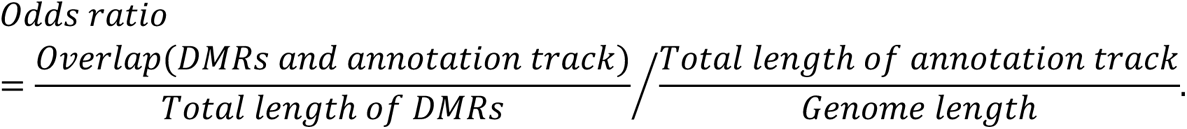

### Gene ontology analysis

We performed GO analysis of tsDMRs shared among COL, PIE and HYB overlapping genes and their ±5 kb neighboring regions, using ShinyGO v0.77 [106] with GO biological process as database. We used an FDR of 0.1 and collared flycatcher as reference annotation.

### Ancestral genome reconstruction and estimates of genetic differentiation

To reconstruct an ancestral genome for the black-and-white flycatchers, we used all-sites genotype data mapped to the COL genome assembly from one red-breasted flycatcher (*Ficedula parva*) and one snowy-browed flycatcher (*F. hyperythra*) previously sequenced [107]. Sites for which red-breasted– and snowy-browed flycatcher were monomorphic were considered callable with the ancestral state being that allele. Sites polymorphic within or between these two species were ignored. To estimate genetic differentiation we used SNPs from 19 COL and 19 PIE flycatchers previously sampled on Öland [95]. Genetic differentiation between COL and PIE was estimated separately for ancestral CpG sites (CpG) and other contexts (non-CpG) using the fixation index *F_ST_* [108] implemented in vcftools v0.1.16 [109].

### Allele-specific methylation estimation

We estimated allele-specific methylation in F1 hybrids using fixed differences (see above) between collared and pied flycatchers as markers within the bisulfite sequencing reads. Allele-specific methylation was called using bismark v0.22.1 *methylation extractor* and the methylation level was measured in 200 bp windows centered at the fixed difference. To measure methylation difference between samples, we used the R package BiSeq v1.28.0 [110]. In F1 hybrids, read coverage were limited to the 0.9 quantile. Parental species for each tissue were randomly downsampled to 3 individuals and coverage was capped at the 0.45 quantile to mimic the sample size and read coverage of HYB. Difference in methylation between groups of samples for each 200 bp locus were determined using the beta regression model in BiSeq v1.28.0 [110]. See Supplementary Methods for more information.

### Statistical framework for molecular mechanism of DNA methylation divergence

We classified the molecular mechanism of DNA methylation divergence by comparing the methylation difference between the parental species and between the parental alleles in the F1 hybrid environment [36, 62]. To ensure that the same number of null hypotheses need to be rejected for calling *cis* and *trans* differences, we also compared the methylation difference of parental alleles and hybrid alleles of the same origin (e.g. PIE: Parental – Hybrid; Table S7). This constitutes a *trans* effect test since by definition we expect the same allele to have different methylation levels when in parent or hybrid environment, if such an effect occurs. Details on the statistical framework are provided in the Supplementary Methods.

## Data access

Bisulfite sequencing data will be available at the European Nucleotide Archive (study ID: PRJEB71458). Scripts are available at the GitHub repository: https://github.com/JesperBoman/DNA_methylation_flycatchers.

## Acknowledgements

The authors thank William Jones, Murielle Ålund, S. Eryn McFarlane, David Wheatcroft, Niclas Backström and Luohao Xu for field and lab work related to this study. The authors are also grateful to Rory J. Craig for contributing to the updated gene annotation and Hans Ellegren, who together with A.Q. and C.F.M. conceived of the study and contributed to the study with the whole-genome bisulfite sequencing data generation. This study was funded by grants from the Swedish Research Council (2013-8271 to Hans Ellegren and 2012-3722 to A.Q.), and the Knut and Alice Wallenberg Foundation (2014/0044 to Hans Ellegren). Sequencing was performed by the SNP&SEQ Technology Platform in Uppsala. The facility is part of the National Genomics Infrastructure (NGI) Sweden and Science for Life Laboratory. The SNP&SEQ Platform is also supported by the Swedish Research Council and the Knut and Alice Wallenberg Foundation. Computations were performed on resources pro-vided by the Swedish National Infrastructure for Computing (SNIC) through Uppsala Multidisciplinary Center for Advanced Computational Science (UPPMAX).

