## Supplementary Information for "Regulatory and evolutionary impact of DNA methylation in two songbird species and their naturally occurring F_1_ hybrids"

1. Supplementary Methods

2. Supplementary Results

3. Figure S1 – Methylation gene profile with gene body split into exons and introns

4. Figure S2 – Tissue-specific patterns of the association between genetic- and methylation differentiation

5. Figure S3 **–** CGI proportion gene profile

6. Figure S4 – Patterns of genetic and epigenetic change at misexpressed genes

7. Table S1 – sample and sequencing information (see external file)

8. Table S2 **–** Frequency of hypermethylation of tissue-specific DMRs

9. Table S3-6 – tsDMR GO analyses (see external files)

10. Table S7 – Classification system for the mechanism of DNA methylation divergence

11. Table S8 **–** Number of fixed difference loci by divergence class

12. Table S9 **–** Number of differentially expressed genes

13. Table S10 **–** Determinants in *cis* of misexpressed genes

Supplementary Methods

**Sampling scheme and tissue collection**

We collected in total 14 male individuals belonging to collared and pied flycatchers, and their naturally occurring F1 hybrids. Initially, four individuals were classified as hybrids using plumage score [1]. Later three of those individuals were confirmed genetically as hybrids (see Mugal et al. 2020 for details), while a fourth was identified as a collared flycatcher (in this study included within the six collared flycatcher individuals).

**Bisulfite sequencing**

DNA was extracted using the phenol-chloroform method. Library preparation and whole-genome bisulfite sequencing were performed by the SciLife SNP&SEQ Technology Platform in Uppsala, Sweden. Sequencing libraries were prepared from 100 ng of DNA using the TruSeq (EpiGnome) Methylation kit (Illumina Inc., EGMK91324) according to the manufacturers’ protocol (#15066014). Samples were multiplexed and split into several lanes as well as separate sequencing runs (Table S1). Sequencing was performed in two separate sequencing efforts, split into a pilot study consisting of brain samples, and a second effort consisting of the remaining samples. Brain samples were sequenced using v4 sequencing chemistry and 125 bp paired-end reads on the Illumina HiSeq2500. In total, 14 lanes were used with two technical replicates and one biological sample per lane (Table S1). The rest of samples were sequenced using v2.5 sequencing chemistry HiSeqX with 150 bp paired-end reads. In total, 87 lanes were used with 6-9 technical replicates per biological sample.

**Processing of bisulfite sequence reads and methylation calls**

Raw reads were quality-controlled using FastQC [3]. Adapter sequences were removed using Trim Galore! v0.6.4_dev, a wrapper for Cutadapt v2.9 [4]. Further clipping was performed according to Epignome profile (8 bp from both 5’ and 3’ ends of both reads in a pair). Reads were aligned to the chromosome version of the collared flycatcher reference genome, FicAlb1.5 [5] using Bismark v0.22.3 with Bowtie2 as alignment tool [6]. Alignment quality was assessed using Qualimap v2.2.2-dev [7] and visualized using MultiQC v1.8 [8]. The percentage of uniquely mapped reads ranged from 50.2 to 70.1 % (Table S1).

**Transcriptome assembly**

Adapters were removed and bases with a Phred score lower than 36 were trimmed from the ends of reads using Trimmomatic v0.39-1 [9]. Reads were further error-corrected using Rcorrector v1.0.3 [10]. Two transcriptome assemblies were created using Spades RNAseq assembler v3.14 with k-mer size 55 and 75 respectively [11]. Further transcriptome assemblies were created with Trans-ABySS v2.0.1 [12] and Trinity v2.9.1 [13]. Assemblies were merged using OrthoFuse [14]. First the separate assemblies were concatenated and groups of transcripts were identified using a modified version of OrthoFinder [15]. The best transcript in each group is then found based on contig score using a modified version of TransRate [16]. Transcripts with less than 1 TPM in the concatenated set of RNA samples and no hit to the Swissprot database were removed from the consensus assembly. The consensus assembly had a TransRate score of 0.1441 and 96.6 % of all BUSCO v3.0.2 genes using the aves_odb9 lineage dataset [17]. Optimal TransRate score for the assembly was 0.2341 but filtering away lower quality contigs reduced the BUSCO coverage to 94 %. Since we here used the transcriptome assembly as a basis for a gene annotation update, we decided to use the more complete assembly of slightly lower contig quality.

**Gene annotation update**

We configured MAKER to update the gene models from the collared flycatcher Ensembl annotation (v96), by setting this annotation as pred_gff in the maker_opts.ctl file. Collared flycatcher RNA-seq evidence for the Ensembl annotation consisted of 8 adult organs/tissues as well as embryo [18]. As additional evidence in the annotation we used the Oyster-River protocol transcriptome assembly, proteins from the chicken (*Gallus gallus,* Ensembl annotation v98) and zebra finch (*Taeniopygia guttata,* Ensembl annotation v102). We configured MAKER to allow gene models to be built directly from transcripts and protein homology. We also included genes found using a pipeline designed to identify so called “missing genes”, i.e. genes that have proven difficult to annotate in the bird genome because of repetitiveness or extreme base composition such as high GC content [19]. Using candidate proteins (n=2454) from the Chinese softshell turtle (*Pelodiscus sinensis,* Ensembl annotation v98) we found collared flycatcher candidate hits (n=1389) using tBLASTn v2.7.1+ to an earlier version of the transcriptome assembly [20]. Candidates were converted to gff3 file format and included as predictions in MAKER (pred_gff option). We used 20,000 bp as the expected max intron size for evidence alignments and conservatively did not consider single exon transcript evidence when generating annotations. We also used an updated repeat annotation consisting of Aves repeats from RepeatMasker v4.0.7_Perl5.24.1, which was mainly repeats curated from chicken and zebra finch, as well as repeats from collared flycatcher [21], hooded crow [22], blue-capped cordon-bleu [23], paradise crow (*Lycocorax pyrrhopterus,* [24]*,* Huon astrapia (*Astrapia rothschildi*), and paradise riflebird (*Ptiloris paradiseus*) [25]. Including consensus sequences derived from TEs in related species has been shown to improve detection of repeats missed by species-specific repeat libraries [23], and should consequently decrease the risk of including TE genes in the gene annotation.

**RNA sequence read analysis**

Quality of raw reads were assessed with FastQC v0.11.9 [3] and adapters were removed using Trim Galore! v0.6.6, a wrapper for Cutadapt v2.10 [4]. Bases with a Phred score < 20 were removed from the 3’ end of reads. Trimmed reads were aligned to the reference genome (see above) using STAR v2.6.1d [26]. Transcript quantification was performed using Salmon v1.4.0 [27]. Quality control was done using a suite of RSeQC v3.0.1 [28], SAMtools v1.1.0 [29] and visualized with MultiQC v1.9 [8].

**A method for enrichment analysis between two sets of genomic ranges using resampling**

Randomization were performed by picking a chromosome at random, weighted by their length and then randomly placing the end coordinate (*i*) somewhere on the chromosome. The start coordinate was then determined by *i* subtracted by the length of DMR. As a heuristic, if *i* subtracted by the length of DMR, was below 0, *i* was considered the start coordinate. We used a two-tailed null hypothesis with a critical value of 0.05 for the resampling procedure. Tests with an empirical *p-*value < 0.025 we considered underrepresented, which means that a set of DMRs and an annotation set overlap less than expected by chance, resulting in an odds ratio significantly <1. Values with a non-significant *p-*value [0.025, 0.975], typically has an odds ratio ~1. We considered tests with a *p-*value > 0.975 enriched, resulting in an odds ratio significantly >1.

**Tissue-specificity of methylation and expression**

Tissue-specific expression was calculated using the preferential expression measure (PEM , Huminiecki et al. 2003) based on the five tissues represented in the study. Genes with no expression in any of the five tissues were excluded. PEM is a relative measure which ranges from -∞ to 1, with values below and above 0 representing relative under- and overexpression respectively. To check for an association between tissue-specific methylation and expression, we selected tsDMRs overlapping annotated promoters. We then tested for a significant deviation in PEM rank for genes with a tsDMR in a focal tissue using *χ^2^* test of independence (*chisq.test* in R). If there is a relationship between tsDMRs and PEM rank, we expect to see the focal tissue with tsDMR in promoter being either the most or least expressed gene among tissues. For tissues and species combinations (e.g. COL x testis) that rejected the null hypothesis of the first test we also proceeded with a secondary test. We tested for a difference in mean PEM between genes with a promoter tsDMR in a focal tissue versus genes with no promoter tsDMR in any tissue (reference set) using the non-parametric Wilcoxon test. Here, we tested for differences between overexpressed (PEM >0) and underexpressed (PEM <0) genes separately. This test investigates the relative importance of DNA methylation in tissue-specific expression since the reference set consists both of genes with ubiquitous expression, as well as tissue-specific genes controlled by mechanisms other than tissue-specific DNA methylation.

**Hybrid inheritance pattern classification framework**

We classified the methylation patterns of F1 hybrids using a cutoff strategy into the following categories of inheritance pattern: conserved, additive, collared-dominant, pied-dominant and the mismethylation categories overdominant and underdominant. Additive means a HYB methylation level between parentals. Collared-dominant corresponds to a methylation level in HYB close to the value of COL and vice-versa for Pied-dominant. Over- and under-dominant means HYB methylation level above and below the value of both COL and PIE respectively. We used a cutoff strategy to determine inheritance pattern instead of statistical tests because different number of significant tests would be needed for different categories. For example, no significant difference between F1 and both parentals for conserved but significantly greater than both parentals for overdominant [31]. However, a simple cutoff strategy is also biased. Consider that we measure the phenotype *P* of two parental species *A* and *B* and their F1 hybrids *H*. If we assume that the phenotypic value *P* is pairwise independent between *A*, *B* and *H* then,

$$cov\left( P_{H}-P_{A}, P_{H}-P_{B} \right)=var\left( P_{H} \right).$$

Since this variance is almost certainly not zero in all practical cases, applying a cutoff will inflate the relative proportion of over- and underdominance. To deal with this artifact, for each promoter we randomly picked a HYB sample for *P_H_* in *P_H_* – *P_A_* and another for *P_H_* in *P_H_* – *P_B_*_._ In effect, this reduces the correlation in error between X and Y axis. Ideally, with a large sample size, many samples would be picked for each group. Simulations showed that this gives a roughly circular error profile around the origin, regardless of whether the measured phenotype is a uniform- or Poisson random variable (data not shown). We thus defined a circular cutoff of 0.1 to classify promoters as either conserved (<0.1) or not (>0.1). To make further classifications as fair as possible we split the X, Y field into 8 areas defined by slices of π/8 radians.

**Allele-specific methylation estimation**

First, deduplicated .bam files of bisulfite sequence reads produced by bismark v0.22.3 were split according to parent-of-origin allele using SNPsplit v0.3.2 [32]. Fixed differences C and T for forward strand alignments and G and A for reverse strand alignments, were ignored since they are indistinguishable from the bisulfite treatment. Allelic imbalance in the number of allele-specific reads were determined with SNPsplit and the kidney sample for HYB02 was excluded due to extreme allelic imbalance in bisulfite-seq reads but not RNA-seq reads (which were analyzed to determine whether the imbalance of the bisulfite-seq reads in HYB02 kidney was a biological effect or a technical artifact). To ensure a sample size of at least 3, kidney was not considered further in allele-specific analyses.

**Statistical framework for molecular mechanism of DNA methylation divergence**

By comparing both COL and PIE methylation differences in HYB and the *trans* effect test we tested both row and column null hypotheses of a 2 x 2 matrix with COL, PIE, parental (PAR) and HYB as column- and row-names, respectively. Due to the dependence of tests in this approach which is a feature it shares with the original framework [33], column-. and row- tests should ideally be done using different samples, though that would require a sample size of hybrids of at least 6. If any sample group lacked methylation read information for a locus, then that locus was classified as *ambiguous*. We classified a locus as *conserved* if there was no difference between COL and PIE in either the parental or HYB comparison. For a *cis*-only change, the COL and PIE allele at a locus had to be significantly different in the same direction both within HYB and between PAR and no significant *trans* effect. Three different outcomes were possible if both a *cis* and a *trans* effect were acting at a locus: 1) *cis* + *trans,* significant difference between PAR and HYB allele for either COL or PIE but with a *trans* effect in the same direction as the *cis* effect, 2) *cis* x *trans, cis* and *trans* effect in opposing directions, 3) *compensatory,* no significant difference in PAR comparison while significant difference in HYB and a *trans* effect. For *trans-*only, there needed to be a significant difference between parentals but not between alleles within F1 hybrids and a significant *trans* effect.

Supplementary Results

1. **Gene annotation**

To improve annotation of untranslated regions (UTRs) and better predict transcription start sites (TSS) and transcription termination sites (TTS), we updated the gene annotation of the collared reference genome FicAlb1.5. For this purpose, we used 36 RNA-seq samples from six tissues (brain, heart, kidney, liver, testis and spleen) of the six collared flycatcher individuals for the multi-assembly Oyster-River protocol [14] for *de novo* transcriptome assembly as well as MAKER [34]. In total, 14,943 out of 16,576 genes mapped to the collared flycatcher chromosome-level assembly. Of these, 9,597 genes had at least one 5’ UTR (in the following referred to as promoter set) and 8563 also had a 3’ UTR (gene profile set).

2. **Relationship between tsDMRs and tissue-specific expression**

To understand the impact of tsDMRs on gene expression levels we quantified tissue-specific gene expression using the preferential expression measure (PEM; Figure SR1) [30]. We tested for a significant deviation from random rank of PEM for genes with a tsDMR in a certain tissue. If tsDMRs are associated with tissue-specific expression, then we expect to see an excess of genes with tsDMRs in the promoter having either the highest or lowest expression. Only tsDMRs at CGI promoters in testis (COL, PIE and HYB) and brain (HYB) had a significant deviation from random PEM ranks (*χ^2^* test of independence, *p <* 0.05), which means that there is an association between tsDMRs and tissue-specific expression. For these, we tested for a significant difference in PEM between genes with a tsDMR in a certain tissue and genes with no tsDMR in any tissue (reference set). This test answers the question: do genes with a detectable tsDMR in the promoter show greater tissue-specific expression than genes lacking promoter tsDMRs? This forms a test of the relative importance of DNA methylation compared to other unobserved explanatory variables for tissue-specific expression patterns. Testis, which had the highest number of tsDMRs overlapping CGI promoters, showed a significantly higher PEM in overexpressed genes compared to the reference set (Wilcoxon test, *p* < 0.05). This highlights the importance of DNA methylation in conferring testis-specific expression, since the reference set consisted both of genes with ubiquitous expression and genes whose tissue-specificity is controlled by mechanisms other than tissue-specific promoter methylation. Underexpressed genes with tsDMRs did not show lower PEM values, indicating that tsDMRs in promoters are generally associated with tissue-specific overexpression in this tissue.

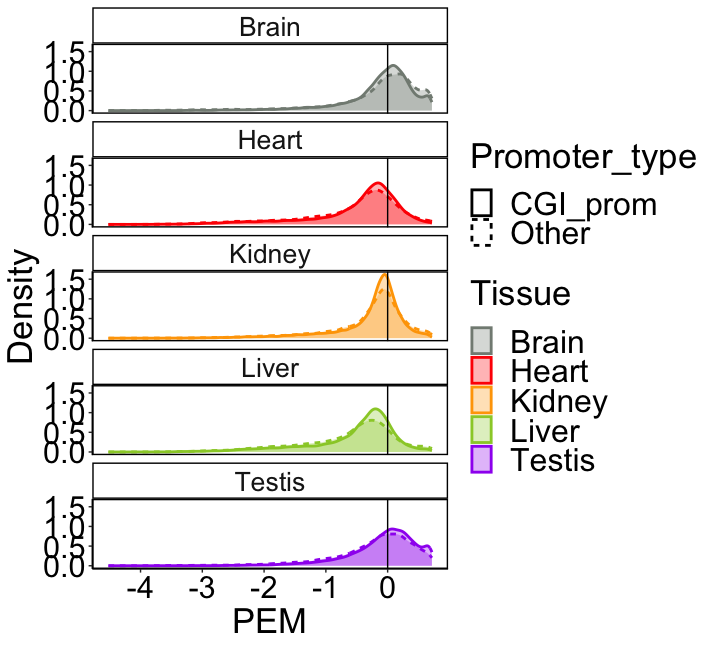

**Figure SR1.** Distribution of tissue-specific expression values per promoter type and tissue. A PEM value >0 and <0 indicates overexpression and underexpression compared to other tissues respectively.

3. **Tissue-specific hybrid inheritance patterns of promoter DNA methylation**

We investigated promoter methylation further because of its role in transcriptional repression. When comparing samples from the parental species using PCA, tissue was a more important factor to divergence in promoter DNA methylation compared to species (Figure SR2A-B). When we split up the dataset by tissue, the PCA separates the species with hybrids grouping intermediately along the first and second PC axes (Figure SR2C-L). This intermediate placement is distinct from earlier observations based on expression data, where misexpression in the hybrids dominated the PCA in all tissues except testis (Mugal et al. 2020).

To further understand the effects of DNA methylation divergence in promoters of hybrids we compared their methylation level with the parental species (see Supplementary Methods). This allowed us to classify the inheritance pattern of DNA methylation into six classes: conserved (HYB close to both COL and PIE), collared-dominant (HYB closer to COL), pied-dominant (HYB closer to PIE), and the two mismethylation categories, overdominant (HYB higher than COL and PIE) and underdominant (HYB lower than COL and PIE). For CGI promoters 81-95 % were classified as conserved per tissue compared to 48-76% for Other promoters (Fisher’s exact test, *p* < 0.05, Figure SR2M). In contrast, the relative proportion of non-conserved inheritance classes were similar between promoter types and only significantly different for heart (*p ≈* 0.033) and liver (*p ≈* 0.013; Figure SR2M). For both heart and liver, more overdominance in *Other* promoters was driving the significance. Around 1-4 % of CGI promoters were mismethylated per tissue compared to 7-11 % for *Other* promoters, further highlighting the stronger conservation of CGI promoters (Figure SR2M). Distribution of inheritance patterns were significantly different between tissues both with (Fisher’s exact test, *p* < 0.05) and without brain, which was sequenced to lower coverage (Table S1). Liver, for example, showed the highest excess of Collared-dominance while heart and testis had excesses of over- and underdominance respectively. These results were in line with genome-wide results indicating that hybrid inheritance of DNA methylation varies predictably among tissues (Table 1). Overall, the intermediate placement of hybrids in-between parental species in the PCA (Figure SR2), is explained by a mixture of collared- and pied- dominant as well as additive effects (Figure SR2M).

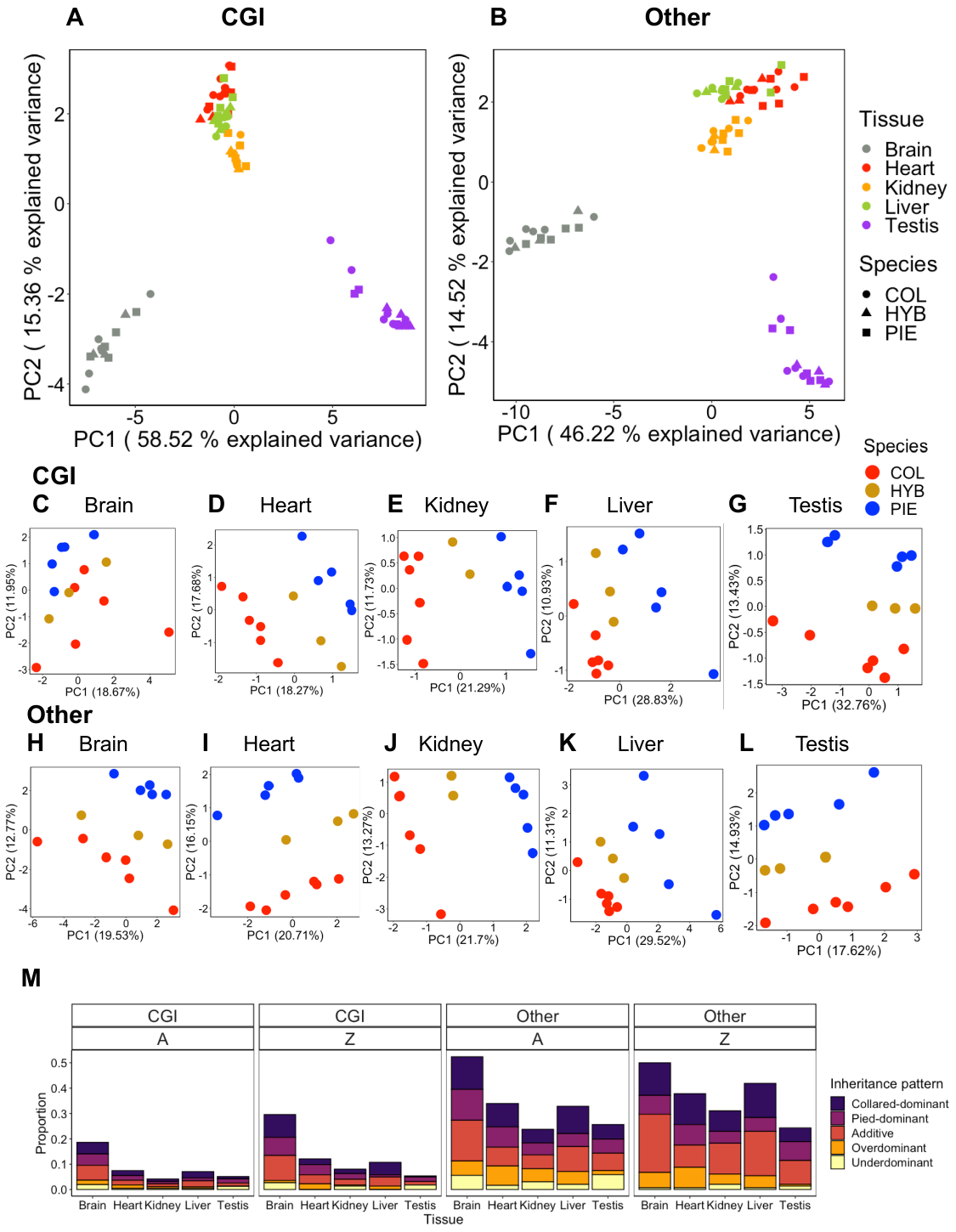

**Figure SR2.** Principal component analysis of promoter methylation patterns and inheritance patterns of methylation in F1 hybrids. CGI promoter methylation PCA for all samples (A) and separated by tissue (C-G). *Other* promoter methylation PCA for all samples (B) and separated by tissue (H-L). When including all samples, tissue dominates methylation variation regardless of promoter type. When separating per tissue, hybrids generally show intermediate promoter methylation levels. The inheritance pattern of promoter methylation shows that this intermediate placement is mainly due to a combination of additive, Collared-dominant and Pied-dominant effects (M).

Previous work on the *Ficedula* flycatchers has shown a greater genetic differentiation on the Z sex chromosome compared to the autosomes (A) [35], as well as divergence in gene expression [2]. While all tissues and promoter type combinations showed a tendency towards less conservation on the Z chromosome, none of these differences were significantly different (Fisher’s exact test, *p >* 0.05). Instead, a slightly higher proportion of additive effects on the Z sex chromosome compared to autosomes where significant for brain (CGI) as well as kidney and liver (*Other*) (Figure SR2M). This may be due to slightly greater methylation differentiation on the Z chromosome (<2 percentage points for all tissues; Wilcoxon rank sum test, *p <* 0.05; Figure SR3), to some extent perhaps caused by greater genetic differentiation (*t-*test; p *<* 2.2 * 10^-16^) on the Z compared to autosomes (Figure SR4). There was also significantly more collared-dominance on Z compared to A in liver and brain (CGI), more pied-dominance on Z for heart (CGI), and less underdominance on Z in brain and testis (*Other*).

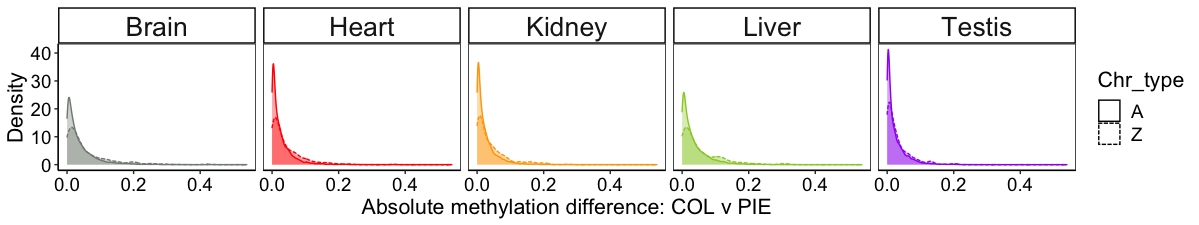
**Figure SR3.** Methylation differentiation (*M_diff_*) of promoter sequences on autosomes (A) and the Z chromosome.

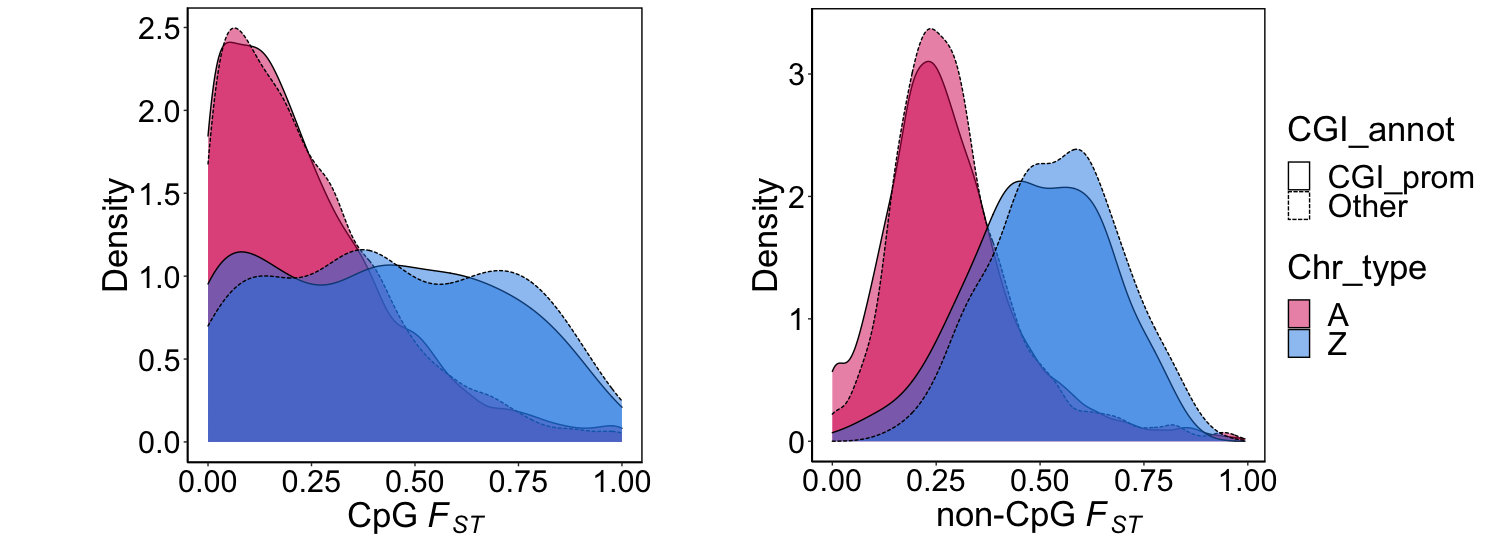

**Figure SR4.** Genetic differentiation (*F_ST_*) in the promoter is higher on the Z compared to the autosomes.

Supplementary Information References

1. Qvarnström A, Rice AM, Ellegren H. Speciation in *Ficedula* flycatchers. Philos Trans R Soc B Biol Sci. 2010;365:1841–52.

2. Mugal CF, Wang M, Backström N, Wheatcroft D, Ålund M, Sémon M, et al. Tissue-specific patterns of regulatory changes underlying gene expression differences among Ficedula flycatchers and their naturally occurring F1 hybrids. Genome Res. 2020;31:1727–39.

3. Andrews S. FastQC: A Quality Control Tool for High Throughput Sequence Data. 2010.

32. Krueger F. SNPsplit v0.3.2. 2017.

33. Fraser HB. Improving Estimates of Compensatory cis–trans Regulatory Divergence. Trends in Genetics. 2019;35:3–5.

34. Holt C, Yandell M. MAKER2: An annotation pipeline and genome-database management tool for second-generation genome projects. BMC Bioinformatics. 2011. https://doi.org/10.1186/1471-2105-12-491.

35. Ellegren H, Smeds L, Burri R, Olason PI, Backström N, Kawakami T, et al. The genomic landscape of species divergence in Ficedula flycatchers. Nature. 2012;491:756–60.

**Figure S1:** Methylation level at CGI (A-E) and *Other* (F-J) promoter types across five tissues with gene body split into intronic and exonic sequence. In general, the methylation level is higher in exons than introns. Orange lines represent intronic sequence and red lines represent exonic sequence. Solid lines represent COL, short-dashed lines HYB and wide-dashed PIE.

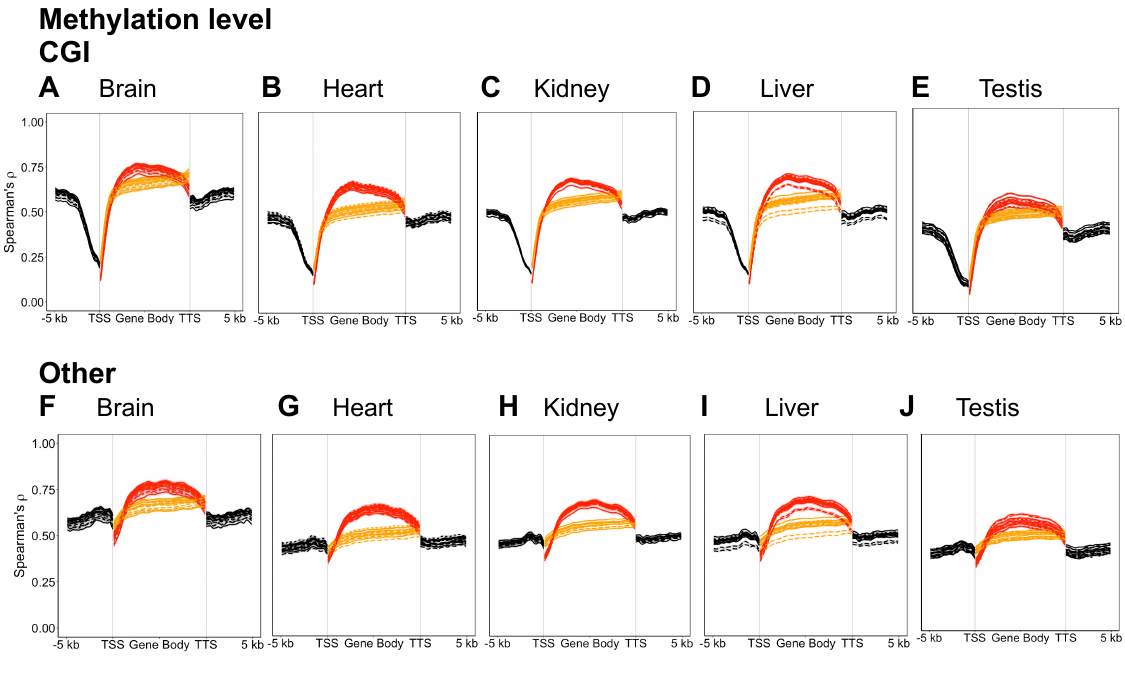

**Figure S2:** Tissue-specific patterns of the association between genetic- and methylation differentiation. (A) and (B) are the same panels as Figure 3A and D of the main text respectively but colored by tissue instead. The results show that tissue both has an impact on the relationship between *M_diff_* and *F_ST_* and that impact is not explained by neither *F_ST_* (which is the same regardless of tissue) nor the average level of *M_diff_*.

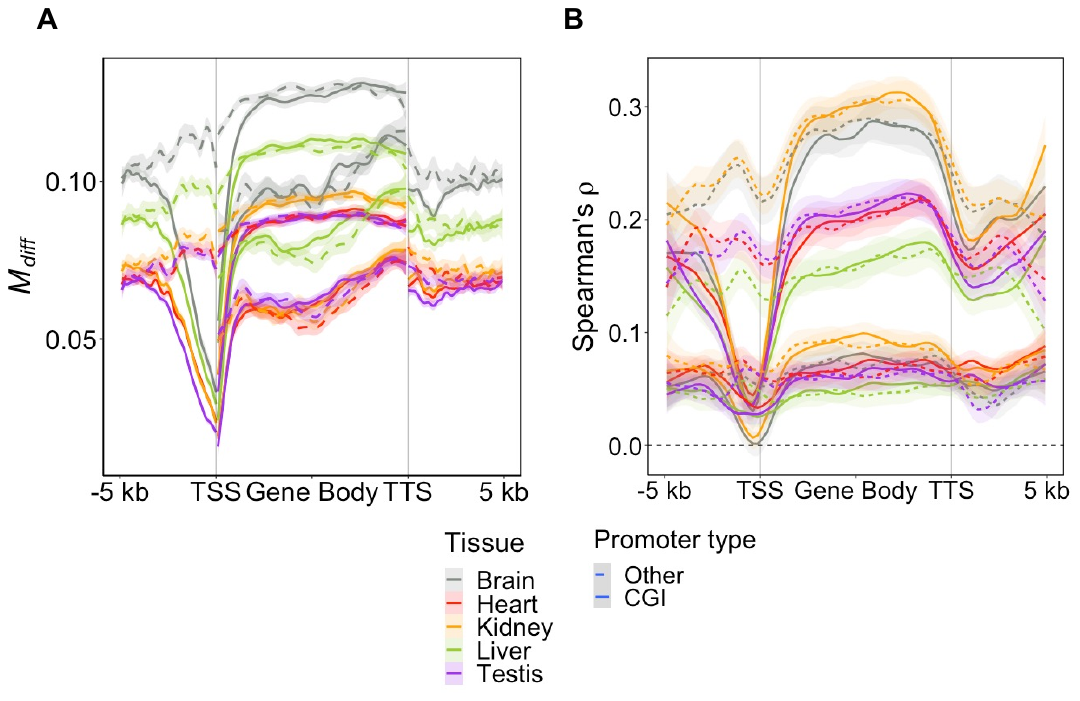

**Figure S3**: Gene profile of the proportion of CGIs. Vertical lines demarcate the TSS and TTS as well as the beginning of the 2kb promoter region.

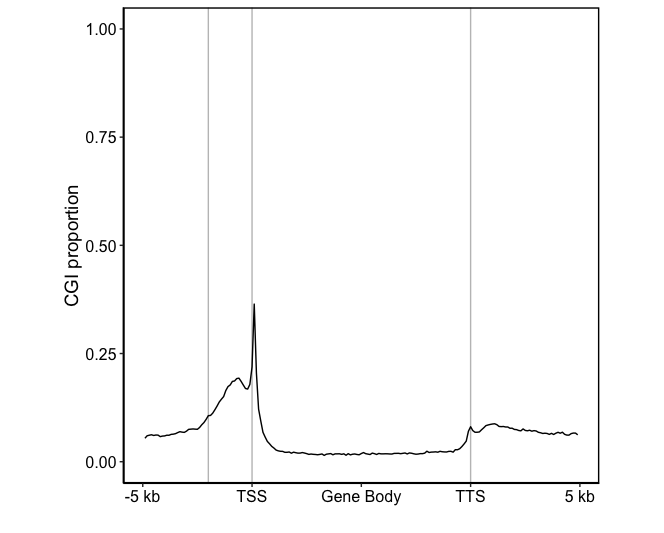

**Figure S4:** Patterns of genetic and epigenetic change at DE genes between HYB and both COL and PIE (misexpressed genes). (A) and (D) show DMR frequency, while (B) and (D) show non-CpG *F_ST_* between COL and PIE. Testis and brain not included in (E) and (F) for visibility due to very few DE genes.

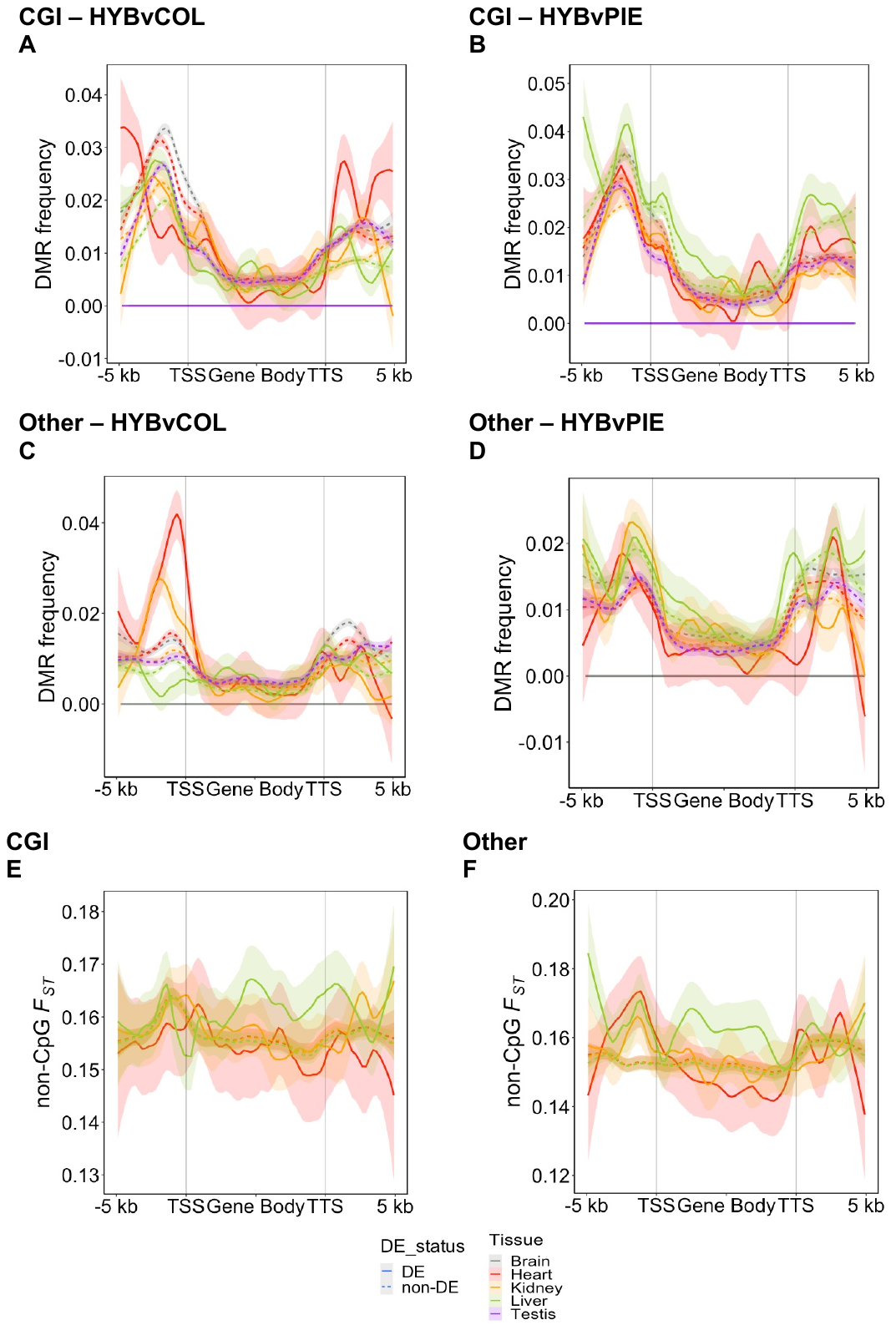

**Table S1**: Sample and sequencing information (see external file).

**Table S2:** Frequency of hypermethylation of tissue-specific DMRs. Here being hypermethylated means that a tsDMR has a higher methylation level in a specific tissue compared to the others. For example, a majority of testis tsDMR were hypermethylated for COL, PIE and HYB.

|  | **Brain** | **Heart** | **Kidney** | **Liver** | **Testis** |
| --- | --- | --- | --- | --- | --- |
| **COL** | 0.61 *** | 0.16 *** | 0.13 *** | 0.33 *** | 0.77 *** |
| **PIE** | 0.56 | 0.13 *** | 0.12 *** | 0.23 | 0.68 *** |
| **HYB** | 0.43 | 0.19 *** | 0.10 *** | 0.17 *** | 0.74 *** |

Family-wise (0.1) adjusted p-value levels * < 0.05 ** < 0.005 *** < 0.0005

**Table S3-6**: tsDMR GO analyses (see external files).

**Table S7:** Classification system for the mechanism of DNA methylation divergence. Classification was based on significance or not in pairwise comparisons between the parental (PAR) species, between alleles in the hybrids (HYB) and between the same allele dependent on parental species background (COL: PARvHYB and PIE: PARvHYB).

| **PAR: COLvPIE** | **HYB: COLvPIE** | **COL: PARvHYB** | **PIE: PARvHYB** | **Results** |
| --- | --- | --- | --- | --- |
| **N** | **N** | **N** | **N** | Conserved |
| **N** | **N** | **S** | **N** | Conserved |
| **N** | **N** | **N** | **S** | Conserved |
| **N** | **N** | **S** | **S** | Conserved |
| **S** | **S** | **N** | **N** | Cis |
| **S** | **S** | **S** | **N** | Cis + Trans or Cis x Trans |
| **S** | **S** | **N** | **S** | Cis + Trans or Cis x Trans |
| **S** | **N** | **S** | **N** | Trans |
| **S** | **N** | **N** | **S** | Trans |
| **S** | **N** | **S** | **S** | Trans |
| **N** | **S** | **S** | **N** | Compensatory |
| **N** | **S** | **N** | **S** | Compensatory |
| **N** | **S** | **S** | **S** | Compensatory |
| **S** | **S** | **S** | **S** | Cis + Trans or Cis x Trans |

**S =** Significant

**N** = Non-significant

**Table S8:** Classifications of fixed difference loci according to methylation patterns in parental species and hybrids. Most loci did not fit the stringent filtering criteria (see main text and Supplementary Methods) and were classified as ambiguous. A majority of loci passing the filtering criteria showed conserved methylation patterns between COL and PIE.

| **Chr_type** | **Tissue** | **Ambiguous** | **Conserved** | **Cis** | **Trans** | **Compensatory** | **Cis x Trans** | **Cis + Trans** |
| --- | --- | --- | --- | --- | --- | --- | --- | --- |
| A | Brain | 30104 | 2916 | 2 | 16 | 4 | 0 | 0 |
| A | Heart | 27556 | 5247 | 20 | 32 | 20 | 6 | 0 |
| A | Liver | 27308 | 5377 | 30 | 36 | 21 | 7 | 1 |
| A | Testis | 25998 | 6714 | 30 | 29 | 41 | 2 | 0 |
| Z | Brain | 3823 | 309 | 1 | 1 | 2 | 0 | 0 |
| Z | Heart | 3496 | 602 | 6 | 2 | 1 | 0 | 0 |
| Z | Liver | 3449 | 631 | 6 | 6 | 2 | 0 | 1 |
| Z | Testis | 3206 | 858 | 12 | 10 | 4 | 0 | 0 |

**Table S9:** Number of differentially- and non-differentially expressed genes in different comparisons split by promoter type.

|  |  | **COL v PIE** | | **HYB v COL** | | **HYB v PIE** | | **Misexpressed** | |
| --- | --- | --- | --- | --- | --- | --- | --- | --- | --- |
| **Tissue** | **Promoter type** | DE | Non-DE | DE | Non-DE | DE | Non-DE | DE | Non-DE |
| Brain | CGI | 52 | 4946 | 1 | 4967 | 9 | 4959 | 1 | 4967 |
| Heart | CGI | 156 | 4842 | 240 | 4727 | 416 | 4551 | 133 | 4834 |
| Kidney | CGI | 308 | 4690 | 533 | 4435 | 1047 | 3921 | 340 | 4628 |
| Liver | CGI | 242 | 4756 | 549 | 4415 | 894 | 4070 | 342 | 4622 |
| Testis | CGI | 687 | 4311 | 24 | 4944 | 44 | 4924 | 1 | 4967 |
| Brain | Other | 29 | 3391 | 6 | 3393 | 5 | 3394 | 1 | 3398 |
| Heart | Other | 84 | 3336 | 162 | 3234 | 263 | 3133 | 95 | 3301 |
| Kidney | Other | 222 | 3198 | 332 | 3067 | 585 | 2814 | 200 | 3199 |
| Liver | Other | 178 | 3242 | 355 | 3043 | 536 | 2862 | 209 | 3189 |
| Testis | Other | 475 | 2945 | 30 | 3369 | 32 | 3367 | 0 | 3399 |

**Table S10:** Determinants in *cis* of overdominant and underdominant genes (misexpressed genes). Average DMR frequency differences and non-CpG *F_ST_* in the 2k upstream promoter region and throughout the gene body were compared between DE and non-DE genes using paired *t-*tests. The table displays p-values of those tests. Brain and testis were excluded since they had too few DE genes.

| **Tissue** | **Promoter type** | **P_DMR (HvP) freq._** | **GB_DMR (HvP) freq._** | **P_DMR_**  **_(HvC) freq_** | **GB_DMR (HvC) freq_** | **P_non-CpG_ *_Fst_*** | **GB_non-CpG_ *_Fst_*** |
| --- | --- | --- | --- | --- | --- | --- | --- |
| Brain | CGI | NA | NA | NA | NA | NA | NA |
| Heart | CGI | 1 (-) | 1 (-) | 0.053 (-) | 0.002 (-) ** | 1 | 1 (-) |
| Kidney | CGI | 1 (-) | 1 (-) | 1 (-) | 0.028 * | 1 (-) | 1 |
| Liver | CGI | 1 | 0.77 | 1 | 1 (-) | 1 (-) | 0 *** |
| Testis | CGI | NA | NA | NA | NA | NA | NA |
| Brain | Other | NA | NA | NA | NA | NA | NA |
| Heart | Other | 1 | 0.124 (-) | 0.001 ** | 1 | 0.321 | 0.011 (-) * |
| Kidney | Other | 0.104 | 1 | 0.087 | 1 (-) | 1 | 1 |
| Liver | Other | 1 | 0.341 | 0.001 (-) * | 0.04 * | 1 | 0 *** |
| Testis | Other | NA | NA | NA | NA | NA | NA |

HvP = Hybrids versus pied flycatchers

HvC = Hybrids versus collared flycatchers

P = Promoter

GB = Gene body

Family-wise (0.1) adjusted p-value levels * < 0.05 ** < 0.005 *** < 0.0005
